# Mu opioid receptors gate the locus coeruleus pain generator

**DOI:** 10.1101/2023.10.20.562785

**Authors:** Makenzie R. Norris, Chao-Cheng Kuo, Samantha S. Dunn, Jenny R. Kim, Léa J. Becker, Gustavo Borges, Loc V. Thang, Kyle E. Parker, Jordan G. McCall

**Affiliations:** Department of Anesthesiology, Washington University in St. Louis, St. Louis, MO, USA; Department of Pharmaceutical and Administrative Sciences, University of Health Sciences and Pharmacy in St. Louis, St. Louis, MO, USA; Center for Clinical Pharmacology, University of Health Sciences and Pharmacy in St. Louis and Washington University School of Medicine, St. Louis, MO, USA; Washington University Pain Center, Washington University in St. Louis, St. Louis, MO, USA; Division of Biology and Biomedical Sciences, Washington University School of Medicine, St. Louis, MO, USA

**Author notes:** To whom correspondence should be addressed: J.G.M.: Address: 660 S. Euclid Ave, Box 8054, St. Louis, MO, USA, 63110. Website: www.mccall-lab.org. These authors contributed equally (Makenzie R. Norris, Chao-Cheng Kuo).

## Abstract

The locus coeruleus (LC) plays a paradoxical role in chronic pain. Although largely known as a potent source of endogenous analgesia, increasing evidence suggests injury can transform the LC into a chronic pain generator. We sought to clarify the role of this system in pain. Here, we show optogenetic inhibition of LC activity is acutely antinociceptive. Following long-term spared nerve injury, the same LC inhibition is analgesic – further supporting its pain generator function. To identify inhibitory substrates that may naturally serve this function, we turned to endogenous LC mu opioid receptors (LC-MOR). These receptors provide powerful LC inhibition and exogenous activation of LC-MOR is antinociceptive. We therefore hypothesized that endogenous LC-MOR-mediated inhibition is critical to how the LC modulates pain. Using cell type-selective conditional knockout and rescue of LC-MOR receptor signaling, we show these receptors bidirectionally regulate thermal and mechanical hyperalgesia – providing a functional gate on the LC pain generator.

## Introduction

The role of the locus coeruleus-noradrenergic (LC-NE) system in chronic pain is both controversial and enigmatic. The LC-NE system is a powerful acute endogenous analgesia system^1^, but evidence also suggests neuropathic injury converts this analgesia system into a system that sustains chronic pain or, in other words, a pain generator^2^. Canonically, the LC modulates descending control of nociception via efferent projections to the dorsal horn of the spinal cord^3^. Supporting this concept, spinal norepinephrine release from LC activation inhibits nociceptive inputs to the dorsal horn via activation of postsynaptic α2 adrenergic receptors^4^. LC-mediated analgesia is further supported by studies showing that cell type- and projection-selective activation of these spinal projections is both antinociceptive and analgesic^1,5^. Recent work, however, demonstrated that activation of LC projections to the deep spinal cord disrupted diffuse noxious inhibitory control of wide dynamic range neurons – a phenomenon thought to underly conditioned pain modulation^6,7^. In this vein, work in human subjects has shown that LC activity is associated with the interaction between attention (increased visual sensory discrimination) and analgesia (diminished nociceptive percept)^8^. The role of the LC in pain becomes further complicated when incorporating studies that disrupt its normal function. In particular, lesioning the LC following neuropathic injury can reverse mechanical hypersensitivity^9,10^ – suggesting some role in maintaining chronic neuropathic pain. While there is evidence that neuropathic injury induces neural plasticity and differential gene expression in LC neurons^11^, it is not immediately clear how these plastic changes convert an analgesic system into a pain generator.

The apparently contradicting role of LC in pain may be due in part to the functional heterogeneity within the LC^12^. Traditionally the LC has been thought of as a mostly homogenous brainstem structure due to its nearly ubiquitous expression of somatic norepinephrine, vast efferent system^13^, and gap junction coupling between LC neurons^14^. However, a growing body of evidence is rapidly redefining this important neuromodulatory system^5,13,15–18^. For example, in rats with neuropathic injury, while chemogenetic activation of LC-NE neurons projecting to the spinal cord expectedly reversed allodynia, activation of LC-NE projections to the medial prefrontal cortex (mPFC) conversely increased spontaneous pain behaviors^5^. Likewise, we recently showed pharmacological inhibition of this mPFC-projecting LC module is antinociceptive^19^. Similarly, extensive work has shown that LC-mediated pain-induced negative affective and cognitive behaviors can be isolated from sensory modulation through distinct projections to the basolateral amygdala, anterior cingulate, hippocampus, dorsal reticular nucleus, and spinal cord^20–23^. While most heterogeneity in the LC has been described in terms of efferent projections, emerging evidence also points to important afferent input to the LC in pain-related behaviors^24–28^.

This afferent control of the LC is mediated through various cell-surface receptors. To that end, LC mu opioid receptors (LC-MOR) have also been implicated in changes to nociceptive processing. While it has long been postulated that LC-MOR dampen the stress response in these neurons^29–34^, the endogenous MOR ligand met-enkephalin also induces antinociception when infused in the LC during the tail flick test^35^. Likewise, LC infusion of the highly selective MOR agonist [D-Ala2, N-MePhe4, Gly-ol]-enkephalin (DAMGO) induces thermal antinociception. This DAMGO-mediated antinociception is reduced during persistent inflammatory pain, likely from the downregulation of LC-MOR protein expression^36^. Neuropathic injury has also been shown to decrease expression of *oprm1* (the gene coding for MOR) in the rat LC^37^. Together, these studies suggest LC-MOR may be a critical component to LC control of nociceptive processing. Here, we show that loss of LC inhibition from endogenous mu opioids may underly sustained pain in injured animals. We use inhibitory optogenetics to isolate the role of LC during acute nociception and nociception following chronic nerve injury. Furthermore, using a conditional knockout approach, we show that LC-MOR expression is required for normal nociception, and that restoring either LC-MOR signaling or receptor expression reverses the hypersensitivity that is caused by spared nerve injury in mice. Together, these data suggest that disruption of LC-MOR-mediated inhibitory tone in the LC converts this analgesic system into a pain generator.

## Results

### High tonic LC activity during hot plate test is not necessary for stress-induced antinociception

Noradrenergic neurons within the LC-NE system canonically exhibit three distinct activation profiles: low tonic, high tonic, and phasic activity. These firing profiles function differently in determining behavioral flexibility to various environmental challenges. Low tonic LC discharge (1-2 Hz) is thought to be consistent with an awake state^38,39^, whereas phasic bursts^18,40^ results from distinct sensory stimuli^38,41,42^. Our previous work, and that by others, has shown that high tonic LC activity (3-8 Hz) drives anxiety-related behavior in mice and rats^15,23,43–46^. However, other studies have demonstrated that this same high tonic activation of LC neurons can be either antinociceptive or pronociceptive, depending on dorsal-ventral localization or efferent projection target^1,5^. These observations led us to hypothesize that stress-induced high tonic LC-NE may contribute to stress-induced antinociception. Indeed, we find that the same 30-minute restraint stress that induces anxiety-like behavior^43^ is also strongly antinociceptive in a hot plate test in mice (**Supplementary Fig. 1A&B**). To determine whether stress-induced high tonic LC activity was responsible for this stress-induced antinociception during noxious stimulation, we used an inhibitory optogenetic approach in mice that previously successfully silenced rat LC neurons^47^. To do so, we selectively expressed a soma-targeted anion-conducting channelrhodopsin (stGtACR2) in the LC of mice heterozygously expressing Cre recombinase in place of dopamine beta hydroxylase (*Dbh*^Cre^::LC-stGtACR2). We first validated this approach with *ex vivo* electrophysiological recordings of locus coeruleus neurons (**Fig. 1A**). Here we show 470 nm activation of stGtACR2 efficiently suppresses spontaneous LC action potentials for up to 35 seconds, substantially longer than the 30-second cutoff we use to prevent tissue damage on the hot plate (**Fig. 1B&C; Supplementary Fig. 2A**). This photoinhibition was particularly efficient, with as little as 2 mW light providing complete inhibition of spontaneous LC firing (**Fig. 1D**). Furthermore, we show that this stGtACR2-mediated inhibition is sufficient to silence activity evoked from large current injections (**Fig. 1E&F**). These recordings suggest stGtACR2 maintains silencing against substantial excitatory input, likely larger than physiological input to the LC *in vivo* during nociception^48^. As might be expected from prior studies with stGtACR2^47,49–51^, whole-cell and cell-attached recordings of stGtACR2 expressing locus coeruleus neurons show rebound firing immediately after blue light cessation (**Supplementary Fig. 2B&C**). This rebound activity, however, is unlikely to affect our behavioral tests, as behavior is recorded and tested exclusively during LC inhibition. To ensure a smooth transition from the restraint stress to hot plate testing, we outfitted mice with a head-fixation bracket around bilateral fiber optic implants above the LC (**Supplementary Fig. 3A**) and restrained animals by both head and body restraint–this approach yields similar antinociception to restraint with a conical tube alone (**Supplementary Fig. 3B**). With all this information in hand, we next created a new cohort of *Dbh*^Cre^::LC-stGtACR2 and *Dbh*^Cre^::LC-mCherry mice with the combined head-fixation bracket and bilateral fiber optic implants above the LC to test whether LC inhibition during the hot plate test could suppress stress-induced antinociception (**Fig. 1G**). Here, LC inhibition had no effect on either *Dbh*^Cre^::LC-stGtACR2 or *Dbh*^Cre^::LC-mCherry stress groups (**Fig. 1H**), suggesting that high tonic LC activity is not necessary for the stress-induced antinociception during noxious sensation. Instead, much to our surprise, LC inhibition in *Dbh*^Cre^::LC-stGtACR2 no stress controls was significantly antinociceptive compared to *Dbh*^Cre^::LC-mCherry no stress controls (**Fig. 1H**). These findings suggest that, basal LC activity, perhaps moreso than in stress states, is critical to normal nociception in naïve animals. We next sought to determine the extent of this contribution across multiple sensory modalities.

**Figure 1.**
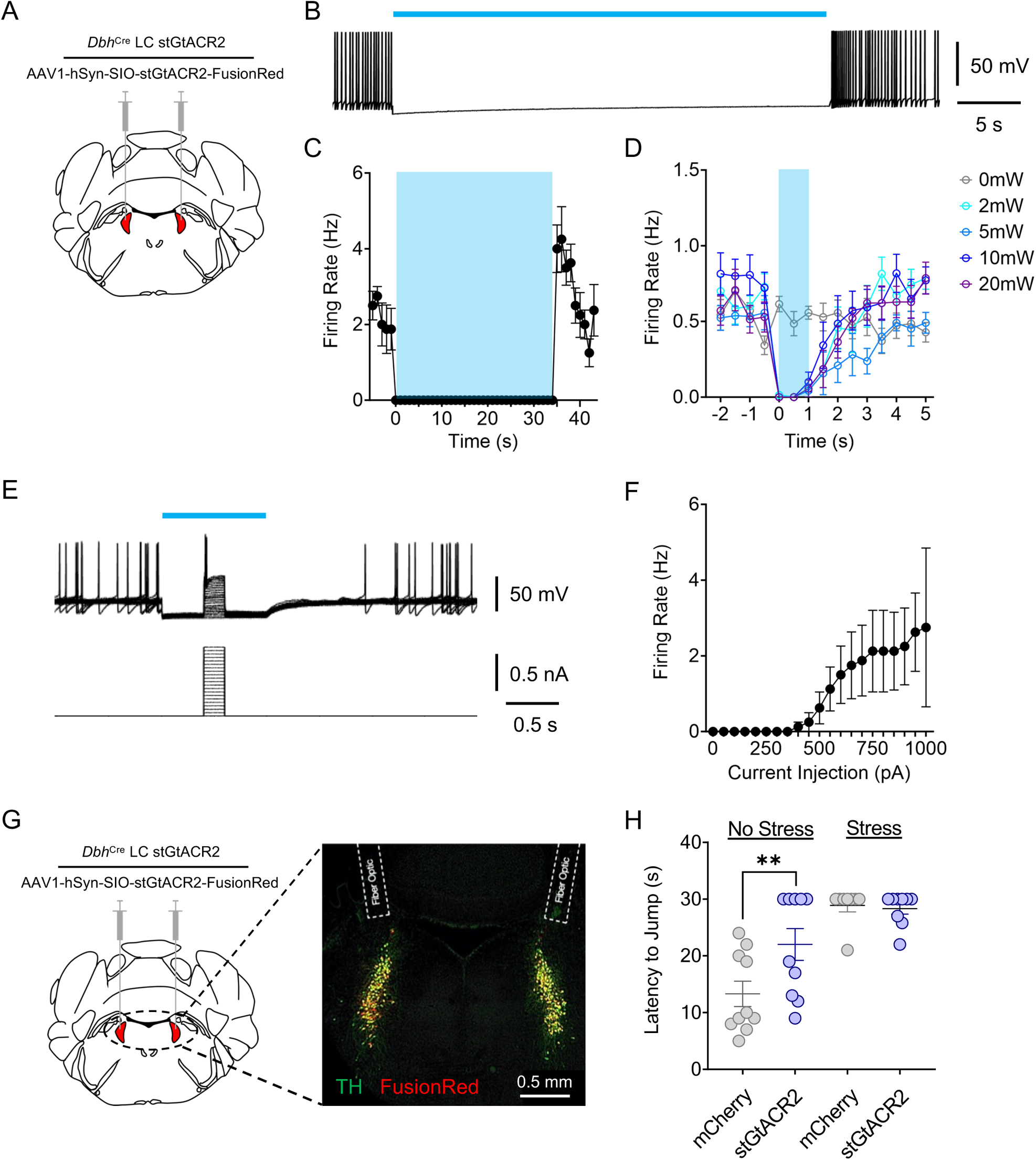
High tonic LC activity during hot plate test is not necessary for stress-induced antinociception. (**A**) Cartoon illustrating the viral strategy for stGtACR2 expression in LC. (**B**) Representative whole-cell recording demonstrating stGtACR2-mediated inhibition of spontaneous firing rate in an LC neuron. (**C**) Quantification of stGtACR2-mediated inhibition of spontaneous LC activity. (**D**) Light power intensity response curve showing 2 mW of 470 nm light illumination is sufficient to suppress the spontaneous activity. (**E**) Representative whole-cell traces demonstrating efficacy of stGtACR2-mediated inhibition of LC neurons against current injections. (**F**) Quantification of stGtACR2-mediated inhibition against current injections. (**G**) Cartoon and fluorescent image of the viral strategy for stGtACR2 expression in LC and bilateral fiber optics implanted above the LC. (**H**) Inhibition of LC neurons following restraint stress does not alter jump latency on a hot plate, but inhibition of LC neurons in stress-naïve mice is antinociceptive. Two-way ANOVA followed by Tukey’s posthoc test, F = 3.901 (stress); 28.13 (photo-Stim.); 5.006 (interaction), ** p<0.01. Data represented as mean ± SEM.

### Intact locus coeruleus activity contributes to baseline hind paw sensory thresholds

While prior studies have shown that high tonic LC activation can be either antinociceptive or pronociceptive^1,5^, these studies did not address the extent to which spontaneous LC activity may contribute to baseline nociceptive thresholds. To determine whether ongoing LC activity is required for baseline nociceptive thresholds in mice, we once again expressed Cre-dependent stGtACR2 or mCherry and implanted fiber optics bilaterally above the locus coeruleus of *Dbh*^Cre^ mice (*Dbh*^Cre^::LC-stGtACR2 or *Dbh*^Cre^::LC-mCherry) (**Fig. 2A**). To determine whether LC activity is required for normal hind paw sensory processing, we inhibited LC neurons during Hargreaves and von Frey tests. LC inhibition increased the thermal stimulus duration necessary to elicit a paw withdrawal in *Dbh*^Cre^::LC-stGtACR2 compared to *Dbh*^Cre^::LC-mCherry, suggesting LC activity is necessary to establish normal evoked thermal thresholds (**Fig. 2B**). Similarly, when we repeated this approach using the Hargreaves test, LC inhibition also significantly increased mechanical withdrawal thresholds (**Fig. 2C**). In these evoked tests, one possible alternative explanation is that LC inhibition produces sedation^39^. However, mice were fully active in the hot plate test, and LC inhibition did not reduce locomotion in a real-time place preference test (**Supplementary Fig. 4**) – in each case showing no signs of sedation. Taken together, these results indicate that spontaneous LC activity helps establish sensory thresholds across both thermal and mechanical domains.

**Figure 2.**
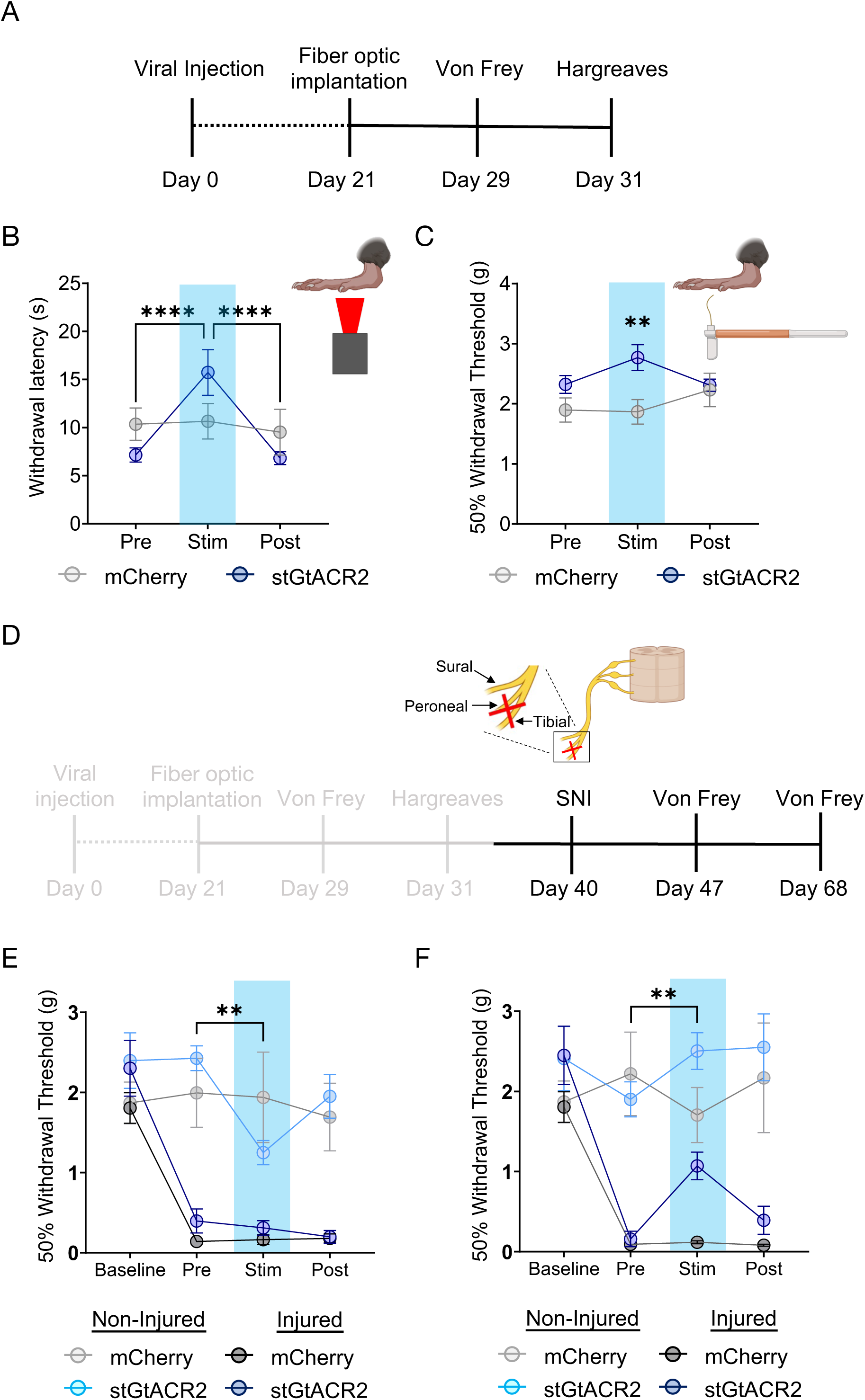
Spontaneous locus coeruleus activity contributes to baseline nociceptive thresholds and adapts over time following injury. (**A**) Experimental timeline. (**B&C**) Hargreaves and von Frey tests showing optical inhibition of LC induces both thermal and mechanical antinociception. Repeated measures two-way ANOVA followed by Tukey’s posthoc test. For thermal nociception (B): F = 10.65 (photo-Stim.); 0.0184 (Inhibition epoch); 7.572 (interaction); 4.290 (animal). For mechanical nociception (C): F = 1.073 (photo-Stim.); 4.922 (Inhibition epoch); 3.861 (interaction); 3.022 (animal),**p<0.01; ****p<0.0001. (**D**) Experimental timeline with gray areas describing data shown in **A-C** and black areas describing data shown in (**E&F**). (**E**) LC inhibition one week after SNI is no longer antinociceptive on the injured limb, but is pronociceptive on the uninjured limb. Repeated measures two-way ANOVA followed by Tukey’s posthoc test, F = 20.05 (SNI development); 20.23 (photo-Stim.); 5.004 (interaction); 1.852 (animal), **p<0.01. (**F**) LC inhibition four weeks after SNI is analgesic at the injured limb. Repeated measures two-way ANOVA followed by Tukey’s posthoc test, F = 9.982 (SNI development); 20.26 (photo-Stim.); 4.416 (interaction); 1.671 (animal), **p<0.01. Data represented as mean ± SEM.

### Locus coeruleus inhibition following neuropathic injury is analgesic in a time-dependent manner

While LC inhibition increases thermal and mechanical thresholds in naïve mice, it is not immediately clear that this same inhibition would provide analgesia in a chronic pain state. To test this hypothesis, we next sought to determine whether the antinociceptive effect of LC inhibition was maintained following neuropathic injury. To do so, we used the spared nerve injury (SNI) model on these same mice, ligating the tibial and peroneal nerve to cause robust and prolonged mechanical hypersensitivity^52–61^(**Fig. 2D**). As expected, one week after SNI, both groups of animals were significantly hypersensitive on the injured limb (**Fig. 2E**). At this one-week timepoint, however, LC inhibition was no longer antinociceptive in the injured limb. Remarkably, this bilateral LC inhibition led to modest, but significant, hypersensitivity in the non-injured limb (**Fig. 2E**). In contrast, four weeks after injury, LC inhibition-induced antinociception was significantly restored in the injured limb of *Dbh*^Cre^::LC-stGtACR2 mice, and the contralateral hypersensitivity in the non-injured limb was no longer present (**Fig. 2F**). This reversal of mechanical hypersensitivity appears to be directly related to noxious sensation, as LC inhibition does not induce a place preference using real-time place testing, providing no evidence of reward from ongoing pain relief (**Supplementary Fig. 4**). These findings further highlight the complexity of LC-mediated analgesia, with both duration and site of injury showing evidence of adaptation over time. Importantly, the analgesic effect of LC inhibition four weeks after SNI is consistent with prior lesion studies that led to the pain generator hypothesis^2,9,10^.

### LC-MOR expression is critical for LC-mediated baseline nociceptive behaviors

Following the observation that exogenous LC inhibition alters nociception in mice, we sought to determine what endogenous inhibitory mechanisms might do the same. MOR are a clear candidate as they are inhibitory G-protein coupled receptors that are heavily expressed in LC neurons and provide potent inhibition when activated^29,62–66^. LC-MOR agonism is antinociceptive, and prior studies have suggested that inflammatory and neuropathic injuries decrease LC-MOR protein and *oprm1* gene expression in the LC^36,37,67^. To determine whether LC-MOR are required for LC-mediated nociceptive control, we generated a noradrenergic neuron-selective conditional knockout mouse through multiple generations of breeding between *Dbh*^Cre+/−^ mice and mice with *loxP* sites on either side of exons 2-3 of the *oprm1* gene (*Oprm1*^fl/fl^)(**Fig. 3A**)^68^. As expected, *Oprm1^fl/fl^xDbh^Cre+/−^* mice present a decrease in *oprm1* mRNA expression in microdissections from the dorsal pons containing the LC (**Fig. 3B**). This knockout strategy appears similar in magnitude to what was previously shown in dorsal root ganglia-selective *oprm1* conditional knockout^68^. We did not observe changes in tyrosine hydroxylase (*th)* expression, the rate-limiting enzyme for NE synthesis, suggesting that the conditional knockout does not disturb catecholamine synthesis (**Fig. 3C**). After reaching *Oprm1*^fl/fl^ homozygosity, *Oprm1*^fl/fl^x*Dbh*^Cre+/−^ LC neurons lose DAMGO-mediated inhibition (**Fig. 3D&E**), consistent with functional disruption of MOR in these cells. Importantly, this LC-MOR deletion does not lead to compensatory changes in spontaneous firing, rheobase, excitability, or input resistance in these cells (**Fig. 3F-I**). We next sought to determine whether *Oprm1*^fl/fl^x*Dbh*^Cre+/−^ mice have normally preserved nociceptive behavioral outputs. To do so, we performed von Frey and Hargreaves testing to determine mechanical and thermal withdrawal thresholds, respectively, in these mice compared to Cre negative controls (*Oprm1*^fl/fl^x*Dbh*^Cre−/−^). We found *Oprm1*^fl/fl^x*Dbh*^Cre+/−^ mice have significantly decreased mechanical and thermal thresholds, suggesting that intact noradrenergic MOR expression is required for normal nociceptive thresholds (**Fig. 3J&K**). This finding is particularly notable given that global MOR deletion (*oprm1^−/−^* mice) decreases thermal nociception and prevents the antinociceptive effects of morphine^69,70^ (**Supplementary Fig. 5**). This difference aligns with the idea that MOR expression outside of noradrenergic cells is critical for typical MOR-mediated analgesia^71–80^. To determine whether removal of any potent endogenous inhibitory system could produce similar effects on nociception, we next generated a noradrenergic neuron-selective conditional knockout mouse of the nociceptin opioid peptide receptor (NOP). NOP and its endogenous ligand nociceptin/orphanin F/Q (N/OFQ) also provide potent inhibition of LC neurons^19^. Taking the same breeding strategy as the noradrenergic MOR conditional knockout (**Supplementary Fig. 6A**), these *Oprl1*^lox/YFP^x*Dbh*^Cre+/−^ mice also lose sensitivity to N/OFQ (**Supplementary Fig. 6B&C**). Importantly, however, nociceptive thresholds are not altered in noradrenergic NOP conditional knockout mice (**Supplementary Fig. 6D&E**) suggesting selectivity of the noradrenergic MOR-mediated changes in nociception. Despite not detecting any compensatory changes in LC electrophysiological properties (**Fig. 3F-I**), *Oprm1^fl/fl^xDbh^Cre+/−^* have LC-MOR deleted throughout development and it is possible that *oprm1* expression is perturbed in other *Dbh*+ cells. To control for these possibilities, we also bilaterally delivered AAV5-hSyn-eGFP-Cre anatomically to the LC of *Oprm1*^fl/fl^ mice (**Supplementary Fig. 7A**). This approach allows for conditional deletion of MOR expression in the LC during adulthood closer to the time of behavioral testing while also maintaining intact MOR expression during development and elsewhere in the body. This viral *Oprm1*^fl/fl^::LC conditional knockout had similar decreased baseline mechanical withdrawal thresholds to the *Oprm1*^fl/fl^x*Dbh*^Cre+/−^ mice (**Supplementary Fig. 7B**). Altogether, deletion of LC-MOR appears to cause a pronociceptive phenotype compared to mice with intact LC *oprm1* expression, suggesting LC-MOR contributes to control of nociception.

**Figure 3.**
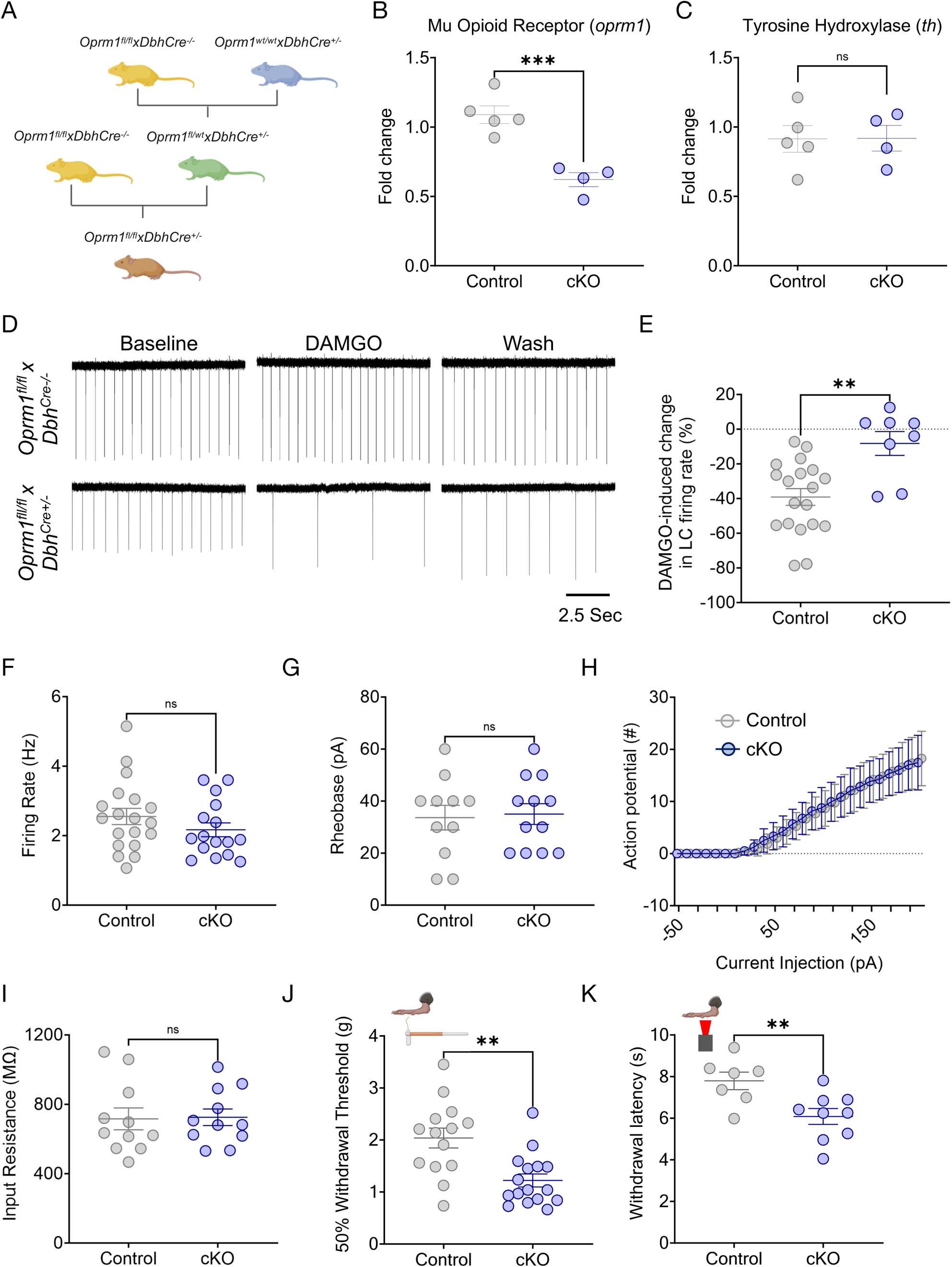
Noradrenergic MOR are required for baseline nociception. (**A**) Schematic describing the breeding strategy of *Oprm1*^fl/fl^x*Dbh*^Cre+/−^ conditional knockout mouse line. (**B&C**) *Oprm1*^fl/fl^x*Dbh*^Cre+/−^ mice show lower expression of *oprm1,* but normal levels of tyrosine hydroxylase (*th*) mRNA in the LC. Student’s t-test. For *oprm1*: t = 5.528 ***p<0.001. For *th*: t = 0.033, ns = not significant. (**D**) Representative cell-attached *ex vivo* LC recordings *Oprm1*^fl/fl^x*Dbh*^Cre−/−^ (top) and *Oprm1*^fl/fl^x*Dbh*^Cre+/−^ (bottom) in response to DAMGO administration. (**E**) DAMGO-mediated inhibition of LC neurons is lost in *Oprm1*^fl/fl^x*Dbh*^Cre+/−^ mice. Mann-Whitney test, U =21, **p<0.01. (**F-I**) Excitability parameters of locus coeruleus neurons between Control (*Oprm1*^fl/fl^x*Dbh*^Cre−/−^) and cKO (*Oprm1*^fl/fl^x*Dbh*^Cre+/−^) mice show no significant differences. These measures include (**F**) baseline firing rate; Mann-Whitney test, U = 116, ns = not significant, (**G**) rheobase; Student’s t-test, t = 0.2218, ns = not significant, (**H**) input-output relationship of number of action potentials fired per current step, and (**I**) input resistance; Student’s t-test, t = 0.1179, ns = not significant. (**J**) von Frey test shows a significant decrease in 50% withdrawal threshold in cKO (*Oprm1*^fl/fl^x*Dbh*^Cre+/−^) compared to Control (*Oprm1*^fl/fl^x*Dbh*^Cre−/−^) mice. Mann-Whitney test, U = 37.5, **p<0.01. (**K**) Baseline thermal withdrawal is also significantly decreased in thermal withdrawal threshold in cKO (*Oprm1*^fl/fl^x*Dbh*^Cre+/−^) mice. Student’s t-test, t = 3.009, **p<0.01. Data represented as mean ± SEM.

### LC-mPFC neurons are crucial for LC-MOR-mediated control of nociception

Given that LC-mediated pain regulation appears to be highly modular^1,5,6,20,23,28^, we suspected that the baseline nociceptive control by LC-MOR could arise from modular functional organization of particular LC efferents. Prior work established that activation of LC projections to the mPFC elicited a pronociceptive tone^5,12^ and facilitates negative affective comorbidities in during neuropathic pain ^22,81^. To determine the basal contribution of this LC-mPFC projection in LC-MOR-mediated control of nociception, we selectively knocked out MOR in LC neurons projecting to the mPFC. Here we bilaterally injected a retrograde canine adenovirus with synthetic promotor targeting noradrenergic neurons (CAV-PRS-Cre-V5) into the mPFC of *Oprm1*^fl/fl^ mice (**Fig. 4A**). The combination of retrograde expression of Cre recombinase and cell type-selective promoter created a module-selective conditional knockout (mKO) of MOR compared to the global conditional knockout of MOR in noradrenergic neurons described above (**Fig. 3**)^82–84^. In a subset of mice, we also used the modular and Cre-dependent expression of mCherry to identify neurons with this genetic and anatomical approach and enable targeted electrophysiological recordings to validate functional deletion. Here DAMGO significantly decreased firing rate in mCherry-expressing LC neurons in C57BL/6J control mice, but not in *Oprm1*^fl/fl^ mice (**Fig. 4A-D**), suggesting successful mKO of MOR in LC-mPFC neurons. We next determined the impact of this mKO strategy on mechanical and thermal nociception. The repeatable nature of these sensory tests allows quantification throughout the genetic development of this mKO. To do so we used von Frey and Hargreaves tests in *Oprm1^fl/fl^*mice receiving either bilateral CAV-PRS-Cre-V5 or CAV-mCherry in the mPFC (**Fig. 4E&F**). Interestingly, we found a progressive within-subject decline of both mechanical and thermal withdrawal thresholds in the CAV-PRS-Cre-V5 group, but not the mCherry controls (**Fig. 4G&H**). Collectively, these results suggest that the MOR in LC-mPFC neurons is critical for MOR-mediated noradrenergic regulation of nociception – a finding in line with the idea that enhanced activity of these neurons promotes nociception and hyperalgesia^5^ and with our contemporaneous results that inhibition of these neurons drive antinociception^19^.

**Figure 4.**
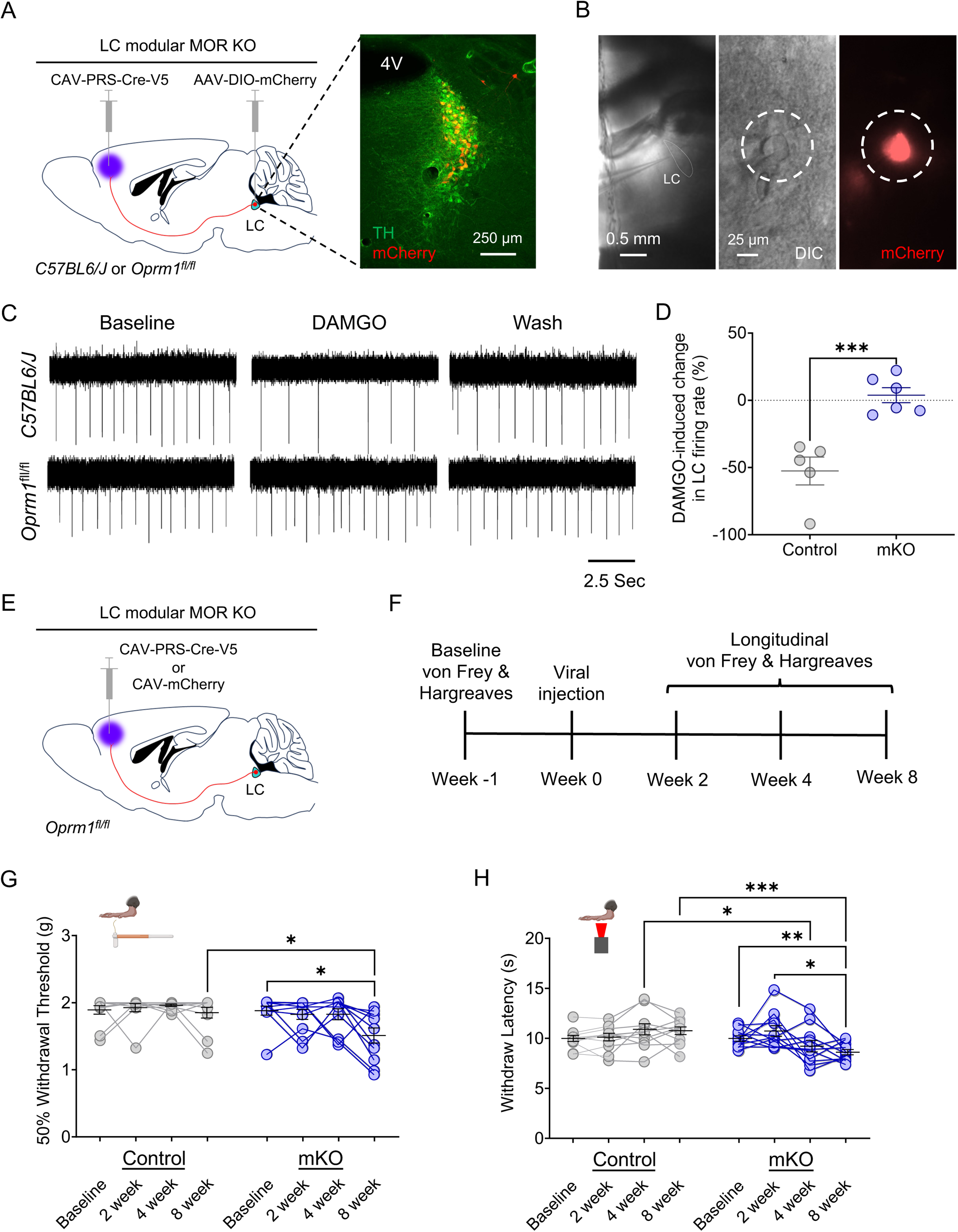
LC-mPFC MOR is crucial for intact LC-mediated antinociception. (**A**) Cartoon and fluorescent image illustrating the viral strategy for modular knockout of MOR in LC for electrophysiological recording. (**B**) Identification of LC-mPFC neurons with module-selective MOR knockout (mKO) by co-expression of mCherry. (**C**) Representative traces of cell-attached recording in LC neurons with an mKO neuron (bottom) showing a loss of DAMGO-mediated inhibition in spontaneous firing rate compared to a C57BL/6J control (top). (**D**) DAMGO-mediated inhibition of LC neurons is lost in mKO-targeted cells of *Oprm1*^fl/fl^ mice. Student’s t-test, t = 5.033 *** p<0.001. (**E**) Schematic illustrating the viral strategy for modular knockout of MOR in LC-mPFC neurons for behavioral testing. (**F**) Timeline of longitudinal nociceptive measurements during the progressive Cre-mediated deletion of modular MOR in LC-mPFC projections. (**G**) von Frey test shows a significant decrease in 50% withdrawal threshold in mKO (*Oprm1*^fl/fl^::mPFC:CAV-PRS-Cre-V5) compared to Control (*Oprm1*^fl/fl^::mPFC:CAV-mCherry) mice. Repeated measures two-way ANOVA followed by Tukey’s posthoc test, F = 5.419 (modular KO); 4.516 (KO progress); 2.057 (interaction); 1.687 (animal), *p<0.05. (**H**) Baseline thermal withdrawal is also significantly decreased in thermal withdrawal threshold in mKO mice. Repeated measures two-way ANOVA followed by Tukey’s posthoc test, F = 3.062 (modular KO); 1.542 (KO progress); 7.140 (interaction); 2.973 (animal), *p<0.05; ** p<0.01; *** p<0.001. Data represented as mean ± SEM.

### Restoration of MOR signaling in the LC reverses Oprm1^fl/fl^xDbh^Cre+^ pronociceptive phenotype

Mice lacking MOR in the LC respond to lower levels of mechanical force and react more rapidly after the application of noxious heat stimuli (**Fig. 3J&K**). To determine whether LC-MOR signaling itself is responsible for this pronociceptive phenotype, we used the light-sensitive chimeric opto-MOR receptor we previously helped develop^85^. Opto-MOR allows for cell type- and intracellular signaling cascade-selective rescue of the G-coupled inhibitory signaling associated with MOR activation. Furthermore, due to its extracellular component unable to bind endogenous MOR ligands, it also leverages temporal control of signaling activation via photostimulation. To test whether this restoration of signaling would restore normal nociception, we used AAV5-Ef1α-DIO-OMOR-EYFP to selectively express opto-MOR in the LC of *Oprm1*^fl/fl^x*Dbh*^Cre+/−^ mice in concert with bilateral fiber optics implantation (**Fig. 5A&B**). Due to the extreme photosensitivity of the chimeric receptor, opaque black caps were used to prevent ambient light from entering the fiber optic ferrules between experiments. We had two different control groups for this experiment. For the first group, we expressed AAV5-Ef1α-DIO-OMOR-EYFP into the LC of *Oprm1*^fl/fl^x*Dbh*^Cre−/−^ mice, as these mice were unable to express the chimeric receptor due to the absence of the Cre recombinase. This group controls for off-target effects of illumination in mice with normal nociception. The second control group expressed AAV5-Ef1α-DIO-EYFP in the LC of *Oprm1*^fl/fl^x*Dbh*^Cre+/−^ mice, as these MOR conditional knockouts maintain the pronociceptive phenotype while also experiencing viral delivery into LC. Opto-MOR activation in the LC of *Oprm1*^fl/fl^x*Dbh*^Cre+/−^ mice increased thermal paw withdrawal latencies in a light-dependent manner with no effect in either control group (**Fig. 5C**). Likewise, opto-MOR LC activation in the same *Oprm1*^fl/fl^x*Dbh*^Cre+/−^ mice also increased mechanical thresholds in the von Frey test (**Fig. 5D**). These findings suggest restoration of LC-MOR signaling alone is sufficient to reverse the pronociceptive phenotype in *Oprm1*^fl/fl^x*Dbh*^Cre+/−^ mice. Similar to when we inhibit LC activity in *Dbh*^Cre+/−^ mice (**Supplementary Fig. 4**), we see no change in locomotion or real-time place preference from opto-MOR activation in *Oprm1*^fl/fl^x*Dbh*^Cre+/−^ mice (**Supplementary Fig. 8**). These data indicate that LC-MOR signaling directly alters thermal and mechanical thresholds.

**Figure 5.**
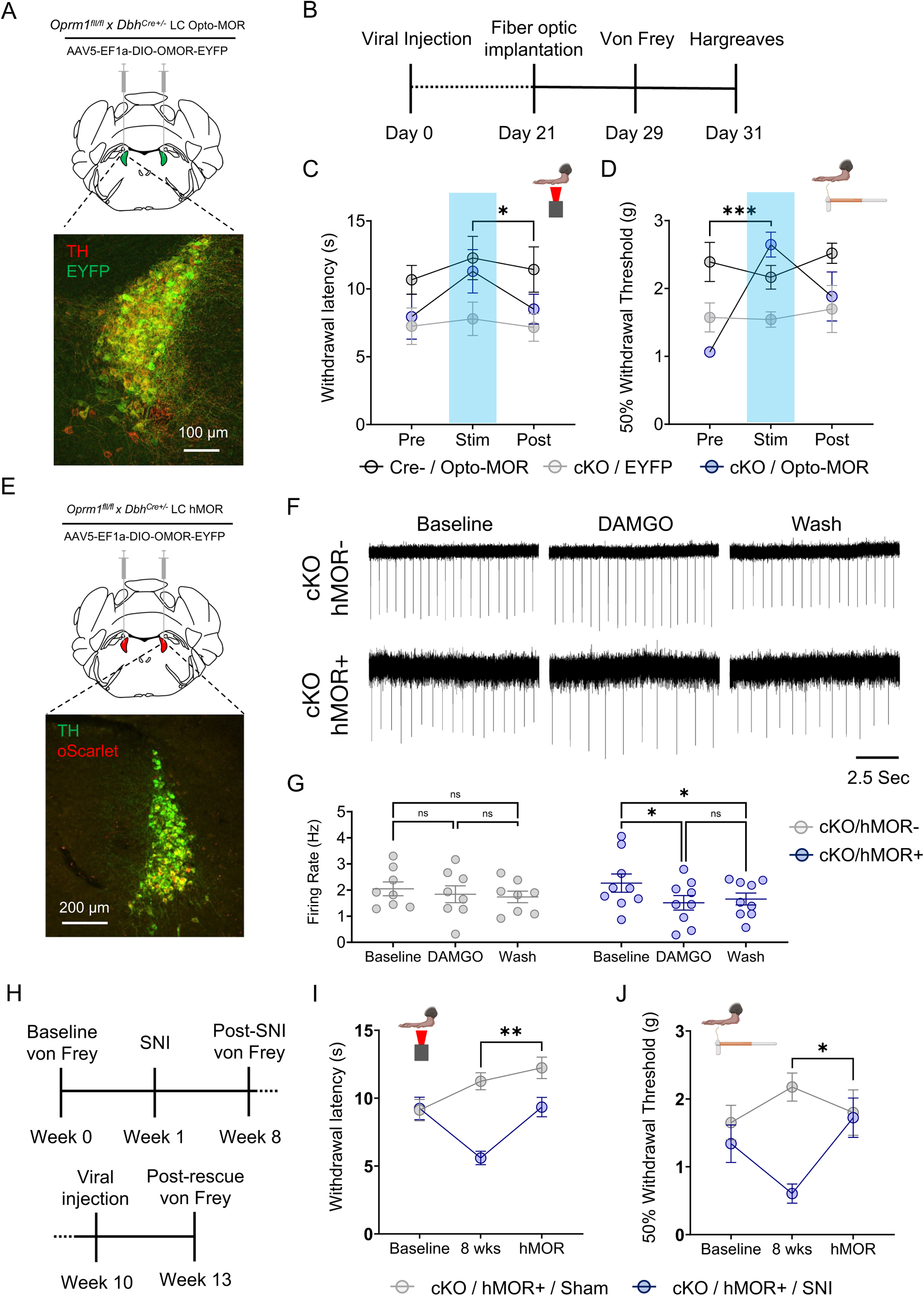
Restoration of LC-MOR signaling reverses baseline hypersensitivity and receptor rescue reverses neuropathic injury-induced hypersensitivity. (**A**) Schematic and fluorescent image for viral strategy expression of opto-MOR in LC. (**B**) Timeline of opto-MOR behavioral tests. (**C**) Opto-MOR activation in the LC restores normal thermal sensitivity. Repeated measures two-way ANOVA followed by Tukey’s posthoc test, F = 3.647 (MOR rescue); 2.764 (Stim. epoch); 0.7294 (interaction); 6.350 (animal), *p<0.05 between cKO:Opto-MOR group Stim vs. Post. (**D**) Opto-MOR activation significantly reverses baseline mechanical hypersensitivity in von Frey test. Repeated measures two-way ANOVA followed by Tukey’s posthoc test, F = 3.749 (MOR rescue); 6.370 (Stim. epoch); 5.554 (interaction); 1.724 (animal), ***p<0.001 between cKO:Opto-MOR Pre vs. Stim groups. (**E**) Schematic and fluorescent image for viral strategy of expression of hMOR in LC. (**F**) Representative traces of cell-attached recordings in LC neurons with hMOR rescued expression show DAMGO-meditated inhibition of LC neurons is restored in *Oprm1*^fl/fl^x*Dbh*^Cre+/−^ mice (bottom) with hMOR compared to those without (top). (**G**) hMOR rescue restores DAMGO-mediated inhibition of LC neurons. Repeated measures two-way ANOVA followed by Tukey’s posthoc test, F = 11.98 (MOR rescue); 0.028 (pharmacology); 3.052 (interaction); 17.47 (cell) **p<0.01. (**H**) Timeline of behavioral measurements following neuropathic injury and rescue of hMOR in the LC. (**I**) hMOR expression reverses neuropathic injury-induced thermal hypersensitivity. Repeated measures two-way ANOVA followed by Tukey’s posthoc test, F = 4.805 (SNI surgery); 34.77 (SNI development & MOR rescue); 6.847 (interaction); 0.558 (animal), **p<0.01. (J) hMOR expression reverses neuropathic injury-induced mechanical hypersensitivity. Repeated measures two-way ANOVA followed by Tukey’s posthoc test, F = 1.034 (SNI surgery); 10.81 (SNI development & MOR rescue); 4.560 (interaction); 0.833 (animal), *p<0.05. Data represented as mean ± SEM.

### Intra-LC MOR rescue in Oprm1^fl/fl^xDbh^Cre+/−^ mice reverses chronic neuropathic injury-induced hypersensitivity

After identifying that LC-mediated baseline nociception is modulated by LC-MOR signaling, we next sought to determine whether full receptor rescue could reverse SNI-induced hypersensitivity. Although *Oprm1*^fl/fl^x*Dbh*^Cre+/−^ mice lack LC-MOR, no modifications were explicitly made to endogenous opioid ligands. If these neuropeptides are still intact in the mice, then rescue of LC-MOR expression should mitigate SNI-induced hypersensitivity. This idea aligns with the previously observed decrease in LC *oprm1* expression after injury, altered opioid-induced activity, and decrease in LC-MOR protein after inflammatory pain^36,37,67^. To test this hypothesis, we first bilaterally expressed the human *OPRM1* gene in the LC of in *Oprm1*^fl/fl^x*Dbh*^Cre+/−^ mice (**Fig. 5E**) and demonstrated that this approach rescues DAMGO-induced LC inhibition in cell-attached recordings (**Fig. 5F&G**). Notably, this inhibition through rescued hMOR expression falls into the physiological range provided by endogenous LC-MOR in the naïve LC (**Fig. 3D&E**), mitigating concern for under- or over-expression in subsequent behavioral experiments. To determine whether endogenous opioid ligands could then restore LC-MOR signaling, we next used a new cohort of *Oprm1*^fl/fl^x*Dbh*^Cre+/−^ mice and performed SNI or Sham surgeries to induce long-term thermal and mechanical hypersensitivity (**Fig. 5H**). Eight weeks after SNI, *Oprm1*^fl/fl^x*Dbh*^Cre+/−^::SNI mice were mechanically and thermally hypersensitive compared to *Oprm1*^fl/fl^x*Dbh*^Cre+/−^::Sham controls (**Fig. 5 I&J**). After establishing this nerve injury-induced hypersensitivity, we then bilaterally expressed human MOR (hMOR) in the LC of both Sham and SNI *Oprm1*^fl/fl^x*Dbh*^Cre+/−^ mice and waited four weeks for expression of the rescued receptor ^86^. Remarkably, we found that rescued hMOR expression in *Oprm1*^fl/fl^x*Dbh*^Cre+/−^::SNI mice completely reversed SNI-induced thermal and mechanical hypersensitivity with no clear effect on *Oprm1*^fl/fl^x*Dbh*^Cre+/−^::Sham controls (**Fig. 5I&J**). These data indicate that rescuing LC-MOR is sufficient for endogenous opioid ligands in the LC to blunt ongoing pain from nerve injury. Altogether, our results suggest LC-MOR-mediated inhibition is critical for evoked sensory responses and, without this LC-MOR-mediated inhibition, the LC generates pronociceptive behavior that can sustain chronic pain.

## Discussion

The locus coeruleus noradrenergic system is well known as a key region in pain neural circuitry^1,3–6,8–11,20,22,87–91^. Several studies have noted that the LC leverages its breadth of efferent circuitry to command robust control over nociceptive processing^1,5–7,20,22,23,28^. Despite elegant previous work, the precise role the LC plays in pain control remains elusive, with clear evidence for both analgesia and generation of chronic pain^1,5,6,9,10,92^. Several studies show the LC mediates acute antinociceptive effects when tonic activity is high^1,5,22,28^. Here we demonstrate that spontaneous LC activity makes important contributions to the evoked behavioral responses from mechanical and noxious thermal stimuli. We first identified this phenomenon while testing the role of LC activity in stress-induced antinociception on the hot plate test (**Fig. 1H**). While this experiment found that LC inhibition during the hot plate test did not alter stress-induced antinociception, further work is necessary to identify whether LC activity during the stressor itself plays a role in restraint stress-induced antinociception. Additionally, we have shown that silencing LC spontaneous activity after nerve injury reveals a time-dependent analgesic effect (**Fig. 2E&F**). While this observation aligns with other studies in rats where LC firing in response to noxious stimuli potentiated under neuropathic pain conditions after four weeks of injury, but not earlier^81^, it also highlights the complexity of the LC’s role in pain because the response to LC inhibition evolves over time and differentially across hindlimbs. One-week after SNI, the injured limb no longer responds to LC inhibition, but the non-injured limb shows enhanced sensitivity. However, four weeks after SNI, LC-mediated inhibition is restored in the injured limb with a trend towards antinociception in the non-injured limb. Following the thread of LC inhibition-mediated control of sensory thresholds, we also report that LC-MOR-mediated inhibition is critical for regulating evoked sensory thresholds (**Fig 3**), while the same is not true for the noradrenergic NOP conditional knockout mice (**Supplementary Fig. 6**). This effect appears to be related to the essential role of MOR expressed in the LC-mPFC efferent module (**Fig 4**). Importantly, in conditional knockout mice lacking MOR in all noradrenergic cells (*Oprm1*^fl/fl^x*Dbh*^Cre+/−^), we found that selectively rescuing LC-MOR signaling or the receptor itself reverses basal pronociceptive phenotypes and SNI-induced hypersensitivity, respectively (**Fig. 5**). These findings add to our understanding of the LC system in pain and suggest that strategies to maintain LC-MOR signaling may be useful for mitigating chronic pain. These results also support the hypothesis that, despite its acute antinociceptive properties, the LC acts as a chronic pain generator^2,6,9,10^. Importantly, our work also adds critical new evidence that loss of LC-MOR may underly the transition from acute analgesia system to chronic pain generator.

This additional insight into the function of LC-MOR expands this receptor system’s functional role in the LC. Most prominently, LC-MOR are thought to bring an end to the LC high tonic stress response^29–34^, but important prior studies identified MOR activation in the LC as antinociceptive^35,36^. Furthermore, long-term neuropathic injury causes acute MOR desensitization and reduced DAMGO responses in rats^93^, and our results in mice suggest this inhibitory G-protein coupled system is critical for regulating LC-mediated nociception. Other contemporaneous work has identified the LC as a critical node for supraspinal exogenous opioid-mediated antinociception through the descending pain system^27^. Altogether, these results suggest rescued mu opioid receptor function may be a therapeutic target for the treatment chronic pain. However, further study is needed to understand how MOR function in LC neurons is altered throughout the duration of neuropathic injury. Although we saw very different results with the noradrenergic NOP conditional knockout mice, it is not yet entirely clear whether MOR function is uniquely required for this effect. Previous studies have shown lesions or blockade of LC activity reverses neuropathic injury-induced hypersensitivity^9,10^. Therefore, it stands to reason that other endogenous inhibitory systems within the LC could produce similar behavioral responses^29,62–66,89,92,94–101^. Our recent work pursued this hypothesis to identify new, potentially analgesic approaches that leverage LC modularity^19^ and more work should be done to identify new modular mechanisms for LC-mediated analgesia.

Altogether, we report here that LC spontaneous activity is critical for normal nociception and regulation chronic nerve injury-induced hypersensitivity. These effects are under endogenous regulation by MOR on LC neurons that project to the prefrontal cortex. These findings have broad implications for our understanding of the chronification of pain and point towards therapeutic solutions. Further study of noradrenergic circuits and inhibitory signaling pathways in these neurons will bring us closer to successfully leveraging the LC system as a target for the treatment of chronic neuropathic pain.

## Materials and Methods

### Animal Subjects

Adult male and female C57BL/6J (JAX:000664), *Dbh*^Cre+/−^ (JAX:033951), *Oprm*1^fl/fl^ (JAX: 030074), *Oprl1*^lox/YFP^ (JAX: 036308)^102^, *Dbh*^Cre+/−^ x *Oprm1^f^*^l/fl^, *Dbh*^Cre−/−^ x *Oprm1^f^*^l/fl^, *Dbh*^Cre+/−^ x *Oprl1* ^lox/YFP^, Dbh^Cre−/−^ x *Oprl1* ^lox/YFP^, and *Oprm1^−/−^* (JAX:007559) mice were used starting from age 8 weeks. Mice were originally sourced from The Jackson Laboratory (Bar Harbor, ME, USA) and bred in-house in a barrier facility in another building. These animals were transferred to a holding facility adjacent to the behavioral space between 4-6 weeks of age. Mice were then left undisturbed except for necessary husbandry to habituate to the new facility until 8 weeks of age. All mice were group-housed, given *ad libitum* access to standard laboratory chow (PicoLab Rodent Diet 20, LabDiet, St. Louis, MO, USA) and water, and maintained on a 12:12-hour light/dark cycle (lights on at 7:00 AM). All experiments and procedures were approved by the Institutional Animal Care and Use Committee of Washington University School of Medicine in accordance with National Institutes of Health guidelines.

### Stereotaxic Surgery

Mice were anaesthetized in an induction chamber (3% isoflurane) and placed in a stereotaxic frame (Kopf Instruments, Model 940) where they were maintained at 1-2% isoflurane. A craniotomy was performed and mice were injected with 250 nl of AAV1-hSyn1-SIO-stGtACR2-FusionRed, AAV1-hSyn-DIO-mCherry, AAV5-syn1-FLEX-oScarlet-T2A-FLAG-hMOR-WPRE, AAV5-Ef1a-DIO-oMOR-eYFP, AAV5-Ef1a-DIO-EYFP, or AAV5-hSyn-eGFP-Cre bilaterally into the LC (stereotaxic coordinates from bregma, anterior posterior (AP): −5.45, medial lateral (ML): +/−1.10, dorsal ventral (DV): −3.75 mm. Mice were then implanted with fiber optic cannula with coordinates adjusted from viral injection −5.45 AP, +/− 1.57 ML, −3.30 DV and implanted at a 10° angle). Implants were secured using Metabond dental cement (C&B Metabond, Edgewood, NY) and super glue. For viral injection into mPFC, 250 nl of CAV-PRS-Cre-V5 or CAV-mCherry was delivered with coordinates as AP: +2.0, ML: 0.4, DV: 1.8 & 0.9. Postoperative care included carprofen tablets and subcutaneous saline injection immediately following surgery. Mice were allowed to have a recovery of 3-6 weeks for AAVs and 7-10 weeks for CAVs prior to behavioral testing and electrophysiological recordings; this interval also permitted optimal viral expression and Cre recombinase activity. pAAV-Syn1-FLEX-mCh-T2A-FLAG-hMOR-WPRE^86^ (Addgene plasmid # 166970; http://n2t.net/addgene:166970; RRID:Addgene_166970) was modified to express the oScarlet fluorophore in place of the original mCherry.

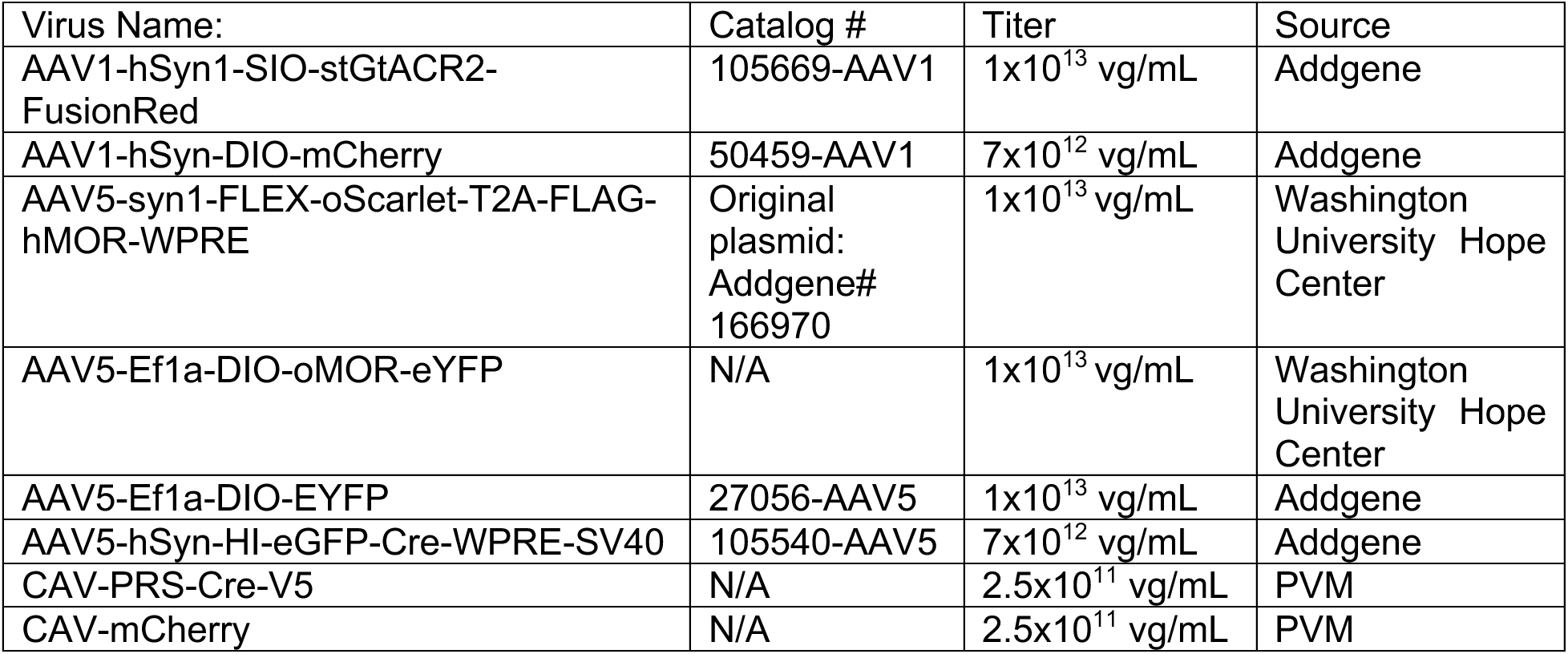

### Stress-Induced Antinociception

Mice were immobilized in modified disposable conical tubes once for 30 minutes as previously described^43^ and were then immediately transferred to the hot plate test.

### Hot plate test

The hot plate apparatus was adapted from the operant thermal plantar assay ^103^. The hot plate was purchased from TE Technologies Inc. (CP-061HT). The Peltier device is independently controlled by a power supply (PS-12–8, 4A, TE Technology) and temperature controller (TC-48– 20, TE Technology). Short cast Acrylic Tubing (7’ height, E-plastics) was used to contain mice on the plate. The plate’s surface temperature was monitored using a surface probe thermometer and maintained at 55°C (Pro-surface thermapen-Thermoworks). Mice were placed onto the hot plate for testing and removed either after 30 seconds of test or after completely a jump defined as both hind paws being removed from the hot plate at once.

### Mechanical sensitivity (von Frey)

Mice were acclimated for 2 hours on an elevated wire mesh grid in 5-inch diameter plexiglass cylinders wrapped in black opaque plastic sheets. Mechanical hypersensitivity was determined by applying von Frey filaments (Bioseb, Pinellas Park, FL,USA) to the lateral aspect of the hind paw using the up-down method as described previously. Von Frey filaments were used at a force ranging from 0.02 g to 3.5 g except for the modular manipulations by CAVs (0.02 to 2g) (**Fig. 4G**). Each von Frey filament stimulation for each mouse was separated by 2 minutes. 50% withdrawal threshold was calculated as previously described^19,60,61^.

### Thermal Plantar Assay (Hargreaves)

Mice were habituated to the Hargreaves apparatus (IITC Life Science, Woodland Hills, CA) in one-hour intervals daily, three days before behavioral testing. On test day, mice were allowed to habituate for 30 minutes before testing. The heat stimulus was set to 40-45% intensity. Paw withdrawal was considered as the paw being removed from the Hargreaves glass surface completely before heat cessation (max duration 30 seconds). For longitudinal measurements of the modular MOR knockout (**Fig. 4H**), intensity of stimulus was adjusted between 20 to 35% to obtain a withdrawal latency around 9 to 12 seconds. Then each mouse was tested with a consistent intensity along the whole measurement. Experimental values were determined by averaging values from left and right foot except where reported separately.

### Spared Nerve Injury (SNI)

The surgical procedure for the SNI-induced model of neuropathic pain was performed as described previously^54,60,61^. Mice were anesthetized with 3% isoflurane and right hind limb shaved and disinfected with 75% ethanol and betadine. A 10-15 mm incision was made in the skin proximal to the knee to expose the biceps femoris muscle. Separation of the muscle allowed visualization of the sciatic nerve trifurcation. The common peroneal and tibial branches were ligated with 6-0 silk suture (Ethicon Inc., Raritan, NJ, USA) and 1 mm of nerve was excised distal to the ligature, leaving the sural branch intact. Following wound closure mice were allowed to recover on a table warmed to 43°C prior to being returned to their home cage. Sham surgeries were identical to the SNI procedure without the ligation, excision, and severing of the peroneal and tibial branches of the sciatic nerve. Behavioral testing on these animals began on post-operative day 7 and wound clips were removed from the healed incision after testing was completed on post-operative day 7. Experimenters were blinded to mouse conditions including sex and injury status during experimental data collection and analysis.

### Real-time place preference test

Animals were placed in a custom-made unbiased, balanced two-compartment conditioning apparatus (52.5 x 25.5 x 25.5 cm). Conditioning apparatus was filled with the same bedding used in mouse home cages equally on both sides. Mice were allowed to freely roam the entire apparatus for 20 min. Entry into one compartment triggered photostimulation of 20 Hz frequency, 2-3mW output for opto-MOR experiments and constant photostimulation, 10 mW output for GtACR2 experiments for the duration the animal remained in the light-paired chamber. Entry into the no light-paired chamber ended photostimulation.

### Immunohistochemistry

Mice were anesthetized with ketamine/xylazine/acepromazine cocktail (69.57 mg/ml; 4.35mg/ml; 0.87mg/ml; i.p. 182mg/kg) and transcardially perfused with ice-cold 4% paraformaldehyde in 1x phosphate buffer saline (PBS). Brains were dissected, post-fixed for 24 hours at 4 °C and cryoprotected with solution of 30% sucrose in 0.1M PB at 4°C for at least 24 hours, cut into 30 µm sections and processed for immunostaining. 30 µm brain sections were washed three times in PBS and blocked in PBS containing 0.5% Triton X-100 and 5% normal goat serum. Sections were then incubated for ∼16 hours at 4°C in rabbit anti-mCherry (600-401-P16, Rockland), chicken anti-TH (1:1000, Aves Labs) or mouse anti-TH (1:500, MilliporeSigma). Following incubation, sections were washed three times in PBS and then incubated for 2 hr at room temperature in Alexa Fluor 594 goat anti-rabbit IgG (1:1000, Invitrogen), Alexa Fluor 488 goat anti-mouse IgG (1:1000, Invitrogen), Alexa Fluor 488 goat anti-chicken IgG (1:1000, Invitrogen), or Alexa Fluor 594 goat anti-chicken IgG (1:1000, Invitrogen) and then washed three times in PBS followed by three 10-min rinses in PB. Sections were then mounted on glass slides with Vectashield Antifade Mounting Medium (Vector Labs) for microscopy. All sections were imaged on an epifluorescent (Leica DM6) or confocal microscope (Leica SP8).

| Antibody name | Host | Target | Catalog | Dilution | Source |
| --- | --- | --- | --- | --- | --- |
| Anti-TH | Chicken | Tyrosine Hydroxylase | TYH | 1:1000 | Aves Labs |
| Anti-TH | Mouse | Tyrosine Hydroxylase | MAB318 | 1:500 | MilliporeSigma |
| Anti-GFP | Chicken | GFP | GP1010 | 1:1000 | Aves Labs |
| Anti-mCherry | Rabbit | mCherry | 600-401-P16 | 1:1000 | Rockland |
| Goat anti-Mouse 488 | Goat | Mouse IgG | A11001 | 1:1000 | Invitrogen |
| Goat anti-Rabbit 594 | Goat | Rabbit IgG | A11012 | 1:1000 | Invitrogen |
| Goat anti-Chicken 488 | Goat | Chicken IgY | A11039 | 1:1000 | Invitrogen |
| Goat anti-Chicken 594 | Goat | Chicken IgY | A11042 | 1:1000 | Invitrogen |

### Electrophysiology

Adult mice were deeply anaesthetized via a ketamine/xylazine/acepromazine cocktail (69.57 mg/ml; 4.35mg/ml; 0.87mg/ml; i.p. 182mg/kg). Upon sedation, mice were perfused with slicing-aCSF consisting of 92 mM N-methyl-d-glucose (NMDG), 2.5 mM KCl, 1.25 mM NaH_2_PO_4_, 10 mM MgSO_4_, 20 mM HEPES, 30 mM NaHCO_3_, 25 mM glucose, 0.5 mM CaCl_2_, 5 mM sodium ascorbate and 3 mM sodium pyruvate, oxygenated with 95% O2 and 5% CO2. pH of aCSF solution was 7.3–7.4 and osmolality adjusted to 315–320 mOsm with sucrose. The brainstem was dissected and embedded with 2% agarose in slice-aCSF. Coronal brain slices were cut into 350μm slices using a vibratome (VF310-0Z, Precisionary Instruments, MA, USA) and incubated in warm (32°C) slicing-aCSF for 30 mins. After incubation slices were transferred to holding-aCSF containing 92 mM NaCl, 2.5 mM KCl, 1.25 mM NaH_2_PO4, 30 mM NaHCO_3_, 20 mM HEPES, 25 mM glucose, 2 mM MgSO_4_, 2 mM CaCl_2_, 5 mM sodium ascorbate and 3 mM sodium pyruvate, oxygenated with 95% O2 and 5% CO2. pH of solution was 7.3–7.4 and osmolality adjusted to 310–315 mOsm. Slices were placed into a recording chamber mounted on an upright microscope (BX51WI, Olympus Optical Co., Ltd, Tokyo, Japan) with epifluorescence equipment and a highspeed camera (ORCA-Flash4.0LT, Hamamatsu Photonics, Shizuoka, Japan) while perfused continuously with warm (29–31°C) recording-aCSF containing 124 mM NaCl, 2.5 mM KCl, 1.25 mM NaH_2_PO_4_, 24 mM NaHCO_3_, 5 mM HEPES, 12.5 mM glucose, 2 mM MgCl_2_, 2 mM CaCl_2_, oxygenated with 95% O2 and 5% CO2 and pH 7.3–7.4 with osmolality adjusted to 305–310 mOsm using sucrose. All recordings were performed using visual guidance (40× water immersion objective lens, LUMPLFLN-40xW, Olympus, Tokyo, Japan) through glass pipette pulled from borosilicate glass capillary (GC150F-10, Warner Instruments, Hamden, CT, USA) with a resistance around 6-9 MΩ. For whole-cell recording, glass pipettes were filled with potassium gluconate-based intra-pipette solution consisting of 120 mM potassium gluconate, 5 mM NaCl, 10 mM HEPES, 1.1 mM EGTA, 15 mM Phosphocreatine, 2 mM ATP and 0.3 mM GTP, pH 7.2–7.3 and osmolality adjusted to 300 mOsm. Data from current-clamp mode were discarded if the membrane potential (Vm) of recorded cell was over −40 mV or action potentials did not overshoot 0 mV. For voltage-clamp recordings, membrane potential was clamped at −70mV and data was only accepted if serial resistance varied smaller than 20% of the baseline value, which was less than 20 MΩ typically. All data were collected using a Multiclamp 700B amplifier (Molecular Devices, San Jose, CA, USA) with a low-pass filtered at 2 kHz and digitized at 10k Hz through Axon Digidata 1440A interface (Molecular Devices, CA, USA) running Clampex software (Molecular Devices, CA, USA). In optogenetic experiments, brain slices were cut from *Dbh*^Cre^ mice bilaterally injected with AAV1-hSyn1-SIO-stGtACR2-FusionRed (Addgene, MA, USA) into the LC and allowed to recover for 5-8 weeks. Blue light pulses were generated from an LED light source delivered through the epifluorescence optical path controlled by Axon Digidata 1440A, light intensity was set to 10mW with 2ms duration at 0.05Hz unless specified otherwise. For input-output relationship and input resistance data, all recordings were performed with synaptic blockers containing 200μM kynurenic acid, 1μM strychnine and 100μM picrotoxin. Recorded cells were continuously clamped at −70mV under current-clamp mode and received one second current injections with 10pA steps. Input resistance was calculated using the linear portion of responses from current injection between −30-0 pA. For pharmacology experiments shown in Fig. 5, *Oprm1*^fl/fl^x*Dbh*^Cre+/−^ mice were bilaterally injected with AAV5-syn1-FLEX-oScarlet-T2A-FLAG-hMOR-WPRE into the LC. Cells were recorded using the cell-attached recording method with pipettes filled with recording-aCSF. Drugs were delivered through the recording-aCSF perfusion system. Electrophysiology data were exported through Clampex software and analyzed using MATLAB (MathWorks, MA, USA) and GraphPad Prism9 (GraphPad Software, MA, USA).

### Tissue collection and RNA extraction

Acute brain slices with 250 μm thickness containing LC region were cut as described for electrophysiology. Bilateral LC were dissected under a microscope (Leica S6E, Leica Microsystem GmbH), immediately frozen and kept at −80 °C until RNA extraction. Total mRNA was extracted from LC tissue using the Arcturus PicoPure RNA Isolation Kit (Thermo Fisher Scientific, Waltham, MA). Around 1 mg of LC tissue was incubated for 30 min in 50 µl of extraction buffer at 42 °C, (500 rpm) and then centrifuged for 2 min at 3,000 g. Supernatant (50 µl) was transferred in a new tube containing 50 µl of 70% EtOH. The mix was then loaded on a RNA Purification Column (pre-conditioned with 250 µl of conditioning buffer for 5 min), centrifuged at 100 g for 2 min (binding of the RNA to the column) and at 16,000 g for 30 s to remove the flowthrough. Next 100 µl of washing buffer 1 (WB1) was added and centrifuged at 8,000 g, for 1min, before adding 10 µl of DNAse and 30 µl of RDD buffer (Qiagen, Germany). The mix was left at room temperature for 15 min before adding 40 µl of WB1 and centrifuge at 8,000 g for 15 s. Column was then washed two times by adding 100 µl of washing buffer 2, centrifuged at 8.000 g for 1 min after the first wash, and two times at 16.000 g for 1 min after the second wash to remove all traces of buffer. Columns were transferred to a new 0.5 ml collection tube, to which 12 µl of elution buffer was added, left at room temperature for 1 min and centrifuge at 1.000 g for 1 min to distribute the elution buffer on the column. Finally, the RNA was eluted by centrifugation for 1 min at 16.000 g. RNA concentration were measured by spectrophotometry (Nanodrop One, Thermo Fisher Scientific, Waltham, MA). Samples were then kept at −80 °C until use.

### RT-qPCR

To generate cDNA, 50 ng of the total mRNA was reversed transcribed with a qScript cDNA synthesis kit (QuantaBio, Beverly, Massachusetts) following manufacturer’s instructions. Real-time quantitative polymerase chain reaction (RT-qPCR) was performed in 10 μL reaction containing 2 μL of cDNA (1/10 dilution), 5 μL of PowerUp SYBR Green Master Mix (Applied Biosystems, Foster City, California, United-States), 2 μL of a mix of forward and reverse primers (10 μM) and 1 μL of H2O. The cycling conditions were 50 °C for 2 min, 95 °C for 10 min, and then 40 cycles at 95 °C for 15 s and 60 °C for 1 min. The following primers were used:

*gadph*: F: TGTCCGTCGTGGATCTGAC; R: CCTGCTTCACCACCTTCTTG

*b2m*: F: TGCTACGTAACACAGTTCCACC; R: TCTGCAGGCGTATGTATCAGTC

*oprm1*: F: CCGAAATGCCAAAATTGTCA; R: GGACCCCTGCCTGTATTTTGT

*th*: F: TGCAGCCCTACCAAGATCAAAC; R: CGCTGGATACGAGAGGCATAGTT

Data were normalized to *gapdh*, *b2m* and *gusb* (from the same animal), and fold changes were calculated using the 2_-ΔΔCt_ method^104^. Dbh^Cre−/−^ x *Oprm1^f^*^l/fl^ mice were used as the control group to normalize the data.

### Statistics and data analysis

All data are expressed as mean ± SEM. In data that were normally distributed, differences between groups were determined using two-tailed independent t-tests, paired-t test, one-way ANOVA, or two-way ANOVA followed by post hoc comparisons if the main effect was significant at p < 0.05. In cases where data failed the Shapiro-Wilk normality test or was ordinal by nature, non-parametric analyses were used. Statistical analyses were conducted using Prism 10.0 (GraphPad). All statistical tests performed as part of this manuscript are available in **Supplemental Datasheet 1**.

## Supporting information

Supplemental Datasheet 1

Supplemental Datasheet 2

## Acknowledgements

We thank the other members of the Al-Hasani and McCall labs for helpful feedback on this project, particularly Manish K. Madasu and Rui-Ni Wu. pAAV-Syn1-FLEX-mCh-T2A-FLAG-hMOR-WPRE^86^ was a gift from Matthew Banghart (Addgene plasmid # 166970; http://n2t.net/addgene:166970; RRID:Addgene_166970). Special thanks to Patricia Jensen for the *Dbh^Cre^*mice, EJ Kremer for the CAVs, Bryan A. Copits for feedback on mouse breeding strategies, and to Daniel C. Castro for the Opto-MOR and hMOR viruses. This work was financially supported by the National Institutes of Health (R01NS117899, J.G.M.; R01NS135401, J.G.M.; F31NS124301, M.R.N.), the McDonnell Center for Systems Neuroscience (J.G.M. and K.E.P.), a Collaboration Support initiative for Translational Anesthesiology Research (COSTAR) award from the Department of Anesthesiology at Washington University School of Medicine (J.G.M.), and the Rita Allen Foundation (J.G.M.) with added financial help from the Open Philanthropy Project (J.G.M.). We would like to acknowledge biorender.com for figure cartoons, the Washington University School of Medicine Hope Center for Neurological Disorders viral vector core, and the Osage Nation, Missouria, Illinois Confederacy, and many other tribes as the ancestral, traditional, and contemporary custodians of the land where this work was conducted.

## Author contributions

M.R.N., C-C. K. and J.G.M. conceived the project and designed the detailed experimental protocols. M.R.N., C.-C.K., S.S.D., J.R.K., L.J.B., G.B., and L.V.T. performed the mouse experiments. M.R.N., C.-C.K., S.S.D., J.R.K., L.J.B., G.B., and L.V.T., and J.G.M performed the investigation and analyzed the data. K.E.P. and J.G.M. conceived and designed the *oprl1* cKO approach. M.R.N., C-C. K., J.R.K. and J.G.M wrote the paper. M.R.N., C-C. K., J.R.K. and J.G.M. edited the paper. M.R.N., K.E.P., and J.G.M. acquired funding. J.G.M. provided research supervision. J.G.M. led overall project administration. All authors discussed the results and contributed to revision of the manuscript.

## Data availability

All data presented in this manuscript is available in **Supplemental Datasheet 2.**

---

**Supplementary Figure 1.**
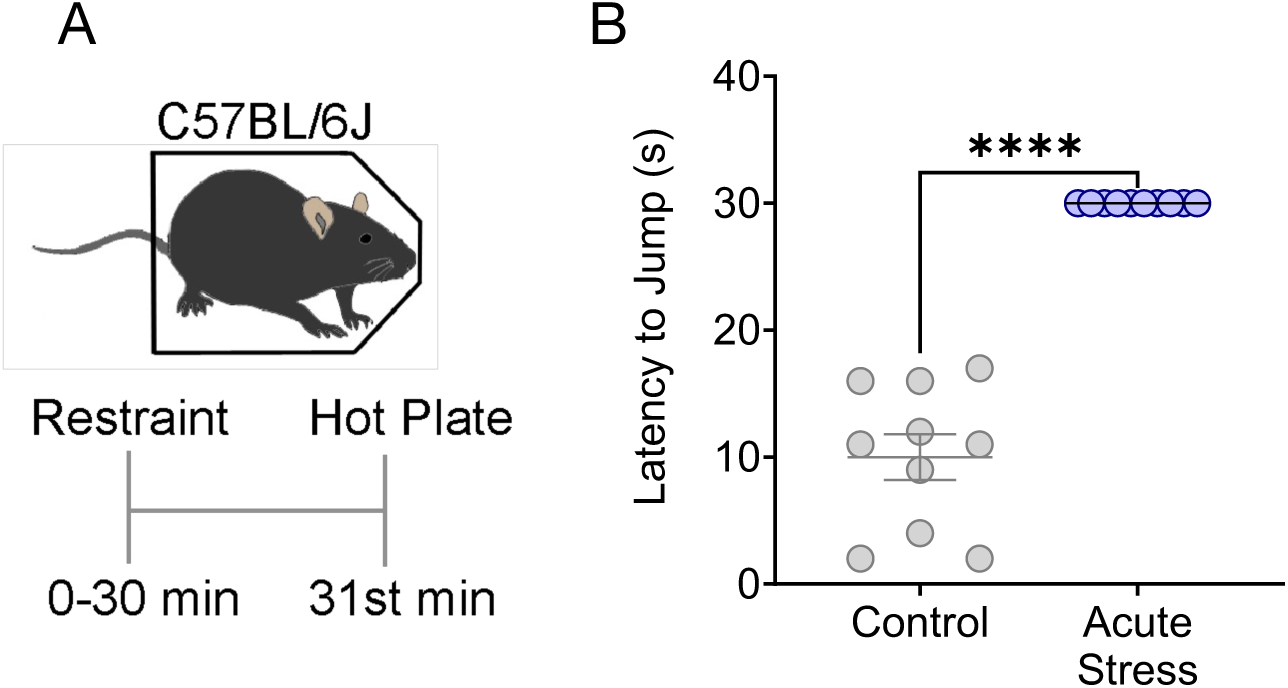
Acute restraint stress causes antinociception. (**A**) Cartoon and experimental timeline for restraint stress-induced antinociception. (**B**) 30 minutes of restraint stress drives thermal antinociception as delayed nocifensive responses on a 55°C hot plate with 30 second cutoff. Student’s t-test, t = 11.1, ****p<0.0001. Data represented as mean ± SEM.

**Supplemental Figure 2.**
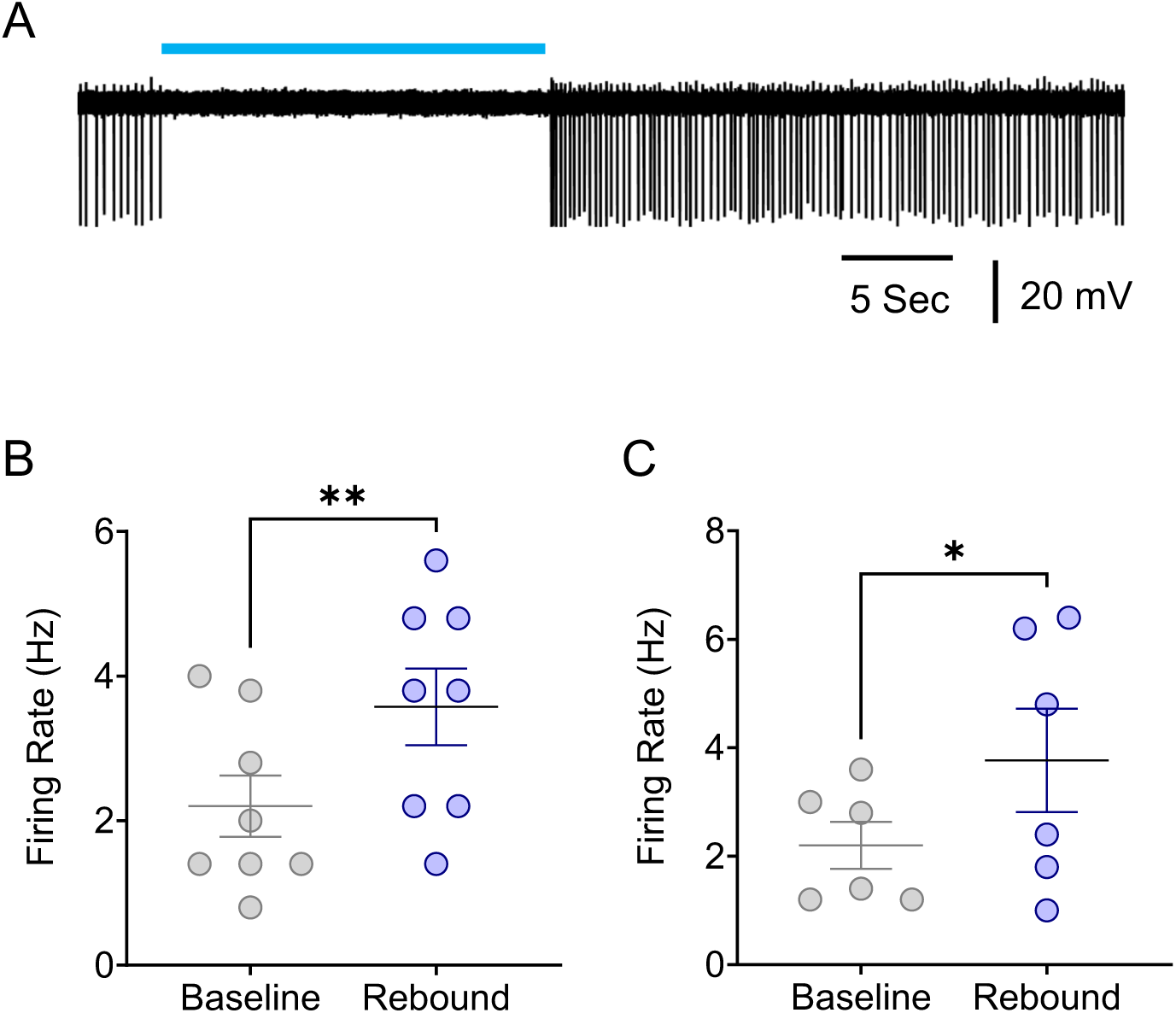
Rebound of neural activity after cessation of stGtACR2-mediated optical inhibition. (**A**) Representative cell-attached recording showing stGtACR2-mediated inhibition in spontaneous firing rate from an LC neuron. (**B&C**) Rebound neural activity following cessation of optical inhibition in either whole-cell (**B**) or cell-attached (**C**) recordings. Paired-t tests, t = 4.604 (whole-cell); 2.731 (cell-attached), *p<0.05; **p<0.01. Data represented as mean ± SEM.

**Supplementary Figure 3:**
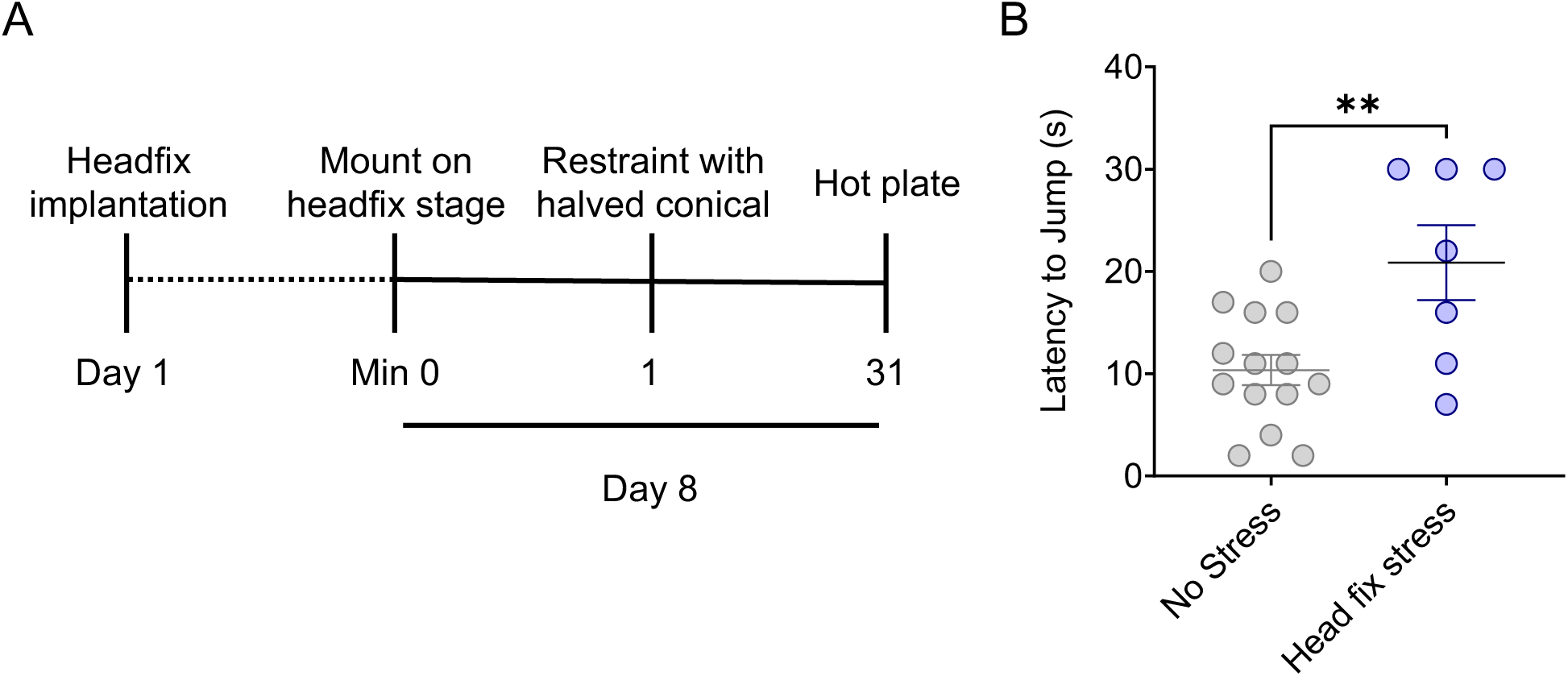
Modified acute restraint stress in concert with head-fixation causes acute antinociception. (**A**) Experimental timeline of modified restraint stress-induced antinociceptive testing in the hot plate test. (**B**) 30 minutes of head-fixed restraint stress drives thermal antinociception as delayed nocifensive responses on a 55°C hot plate with 30 second cutoff. Student’s t-test, t = 3.184, **p<0.01. Data represented as mean ± SEM.

**Supplement Figure 4.**
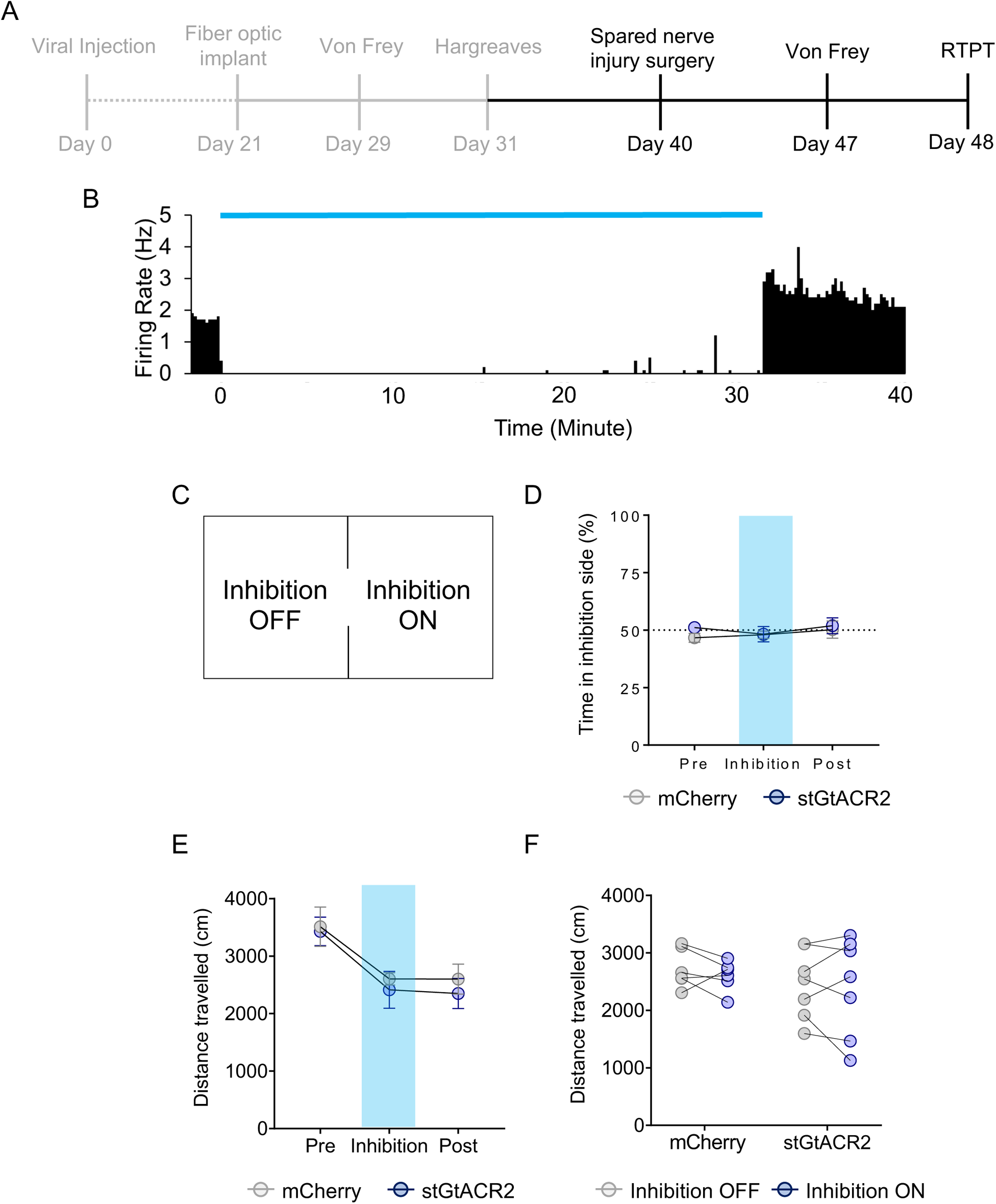
LC optical inhibition does not induce real-time preference four weeks after neuropathic injury. (**A**) Experimental timeline of behavioral testing. (**B**) Histogram showing prolonged LC optical inhibition for 30 minutes using 100 Hz, 0.5 ms pulse width, 470 nm illumination. (**C**) Diagram of real-time place testing apparatus. (**D**) LC inhibition does not induce real-time place preference four weeks post-SNI. Repeated measures two-way ANOVA followed by Tukey’s posthoc test, F = 0.604 (Inhibition epoch); 0.773 (photo-inhibition); 0.302 (interaction); 1.152 (animal). (**E&F**) LC inhibition does not alter distance travelled in real-time place test. Repeated measures two-way ANOVA. For (E): F = 14.41 (inhibition epoch); 0.346 (photo-inhibition); 0.082 (interaction); 2.965 (animal). For (F): F = 0.559 (inhibition On/Off); 0.643 (photo-inhibition); 0.119 (interaction); 8.129 (animal). Data represented as mean ± SEM, no significant difference was found.

**Supplement Figure 5.**
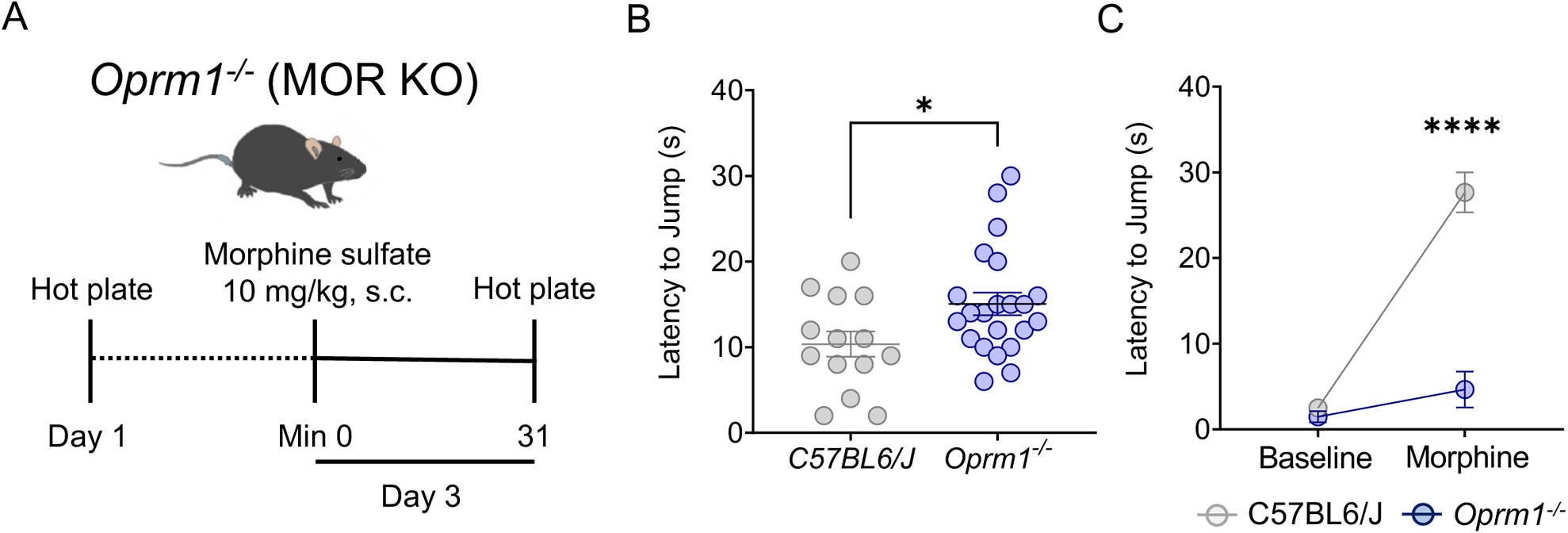
Global *Oprm1* knockout influences nociceptive responses. (**A**) Experimental timeline for behavioral tests. (**B**) Naïve *oprm1*^−/−^ mice have increased nociceptive thresholds on the hot plate. Student’s T-test, t = 2.297, *p<0.05. (**C**) *Oprm1*^−/−^ mice do not have morphine analgesia. Repeated measures two-way ANOVA followed by Tukey’s posthoc test, F = 48.41 (MOR KO); 37.60 (Morphine application); 29.14 (interaction); 0.927 (animal), ****p<0.0001. Data represented as mean ± SEM.

**Supplement Figure 6.**
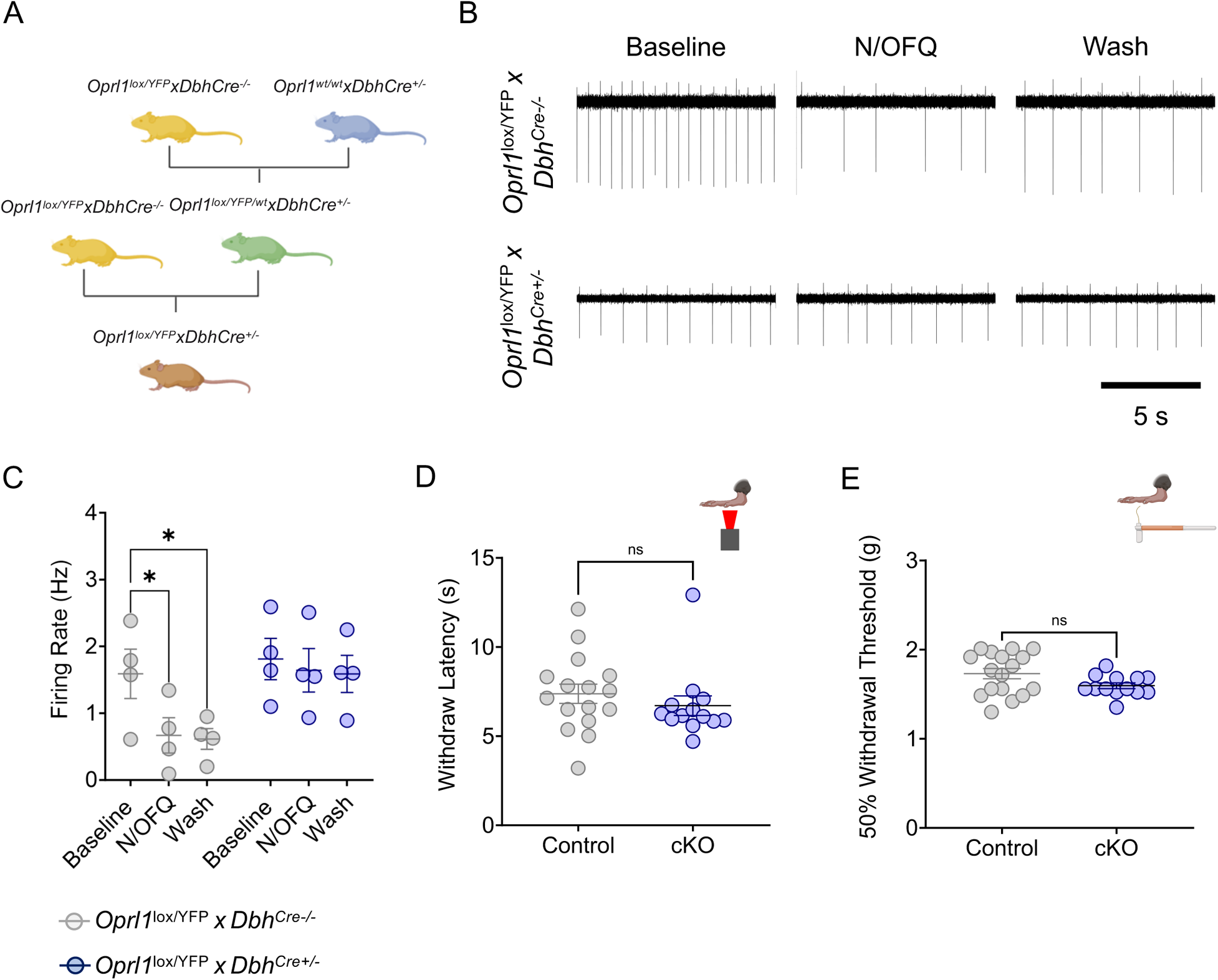
Noradrenergic NOP is not required for noradrenergic-modulated nociception. (**A**) Schematic describing the breeding strategy of *Oprl1*^lox/YFP^x*Dbh*^Cre+/−^ conditional knockout mouse line. (**B**) Representative traces of cell-attached LC recordings with NOP cKO (*Oprl1*^lox/YFP^x*Dbh*^Cre+/−^) losing N/OFQ-mediated LC inhibition (bottom) compared to Control mice (*Oprl1*^lox/YFP^x*Dbh*^Cre−/−^). (**C**) N/OFQ-mediated inhibition of LC neurons is lost in *Oprl1*^lox/YFP^x*Dbh*^Cre+/−^ mice. Repeated-measures two-way ANOVA, F = 24.86 (cKO); 3.306 (pharmacology); 10.68 (interaction); 26.80 (cell), *p<0.05. (**D&E**) Hargreaves (Mann-Whitney, U = 69, ns = not significant) (**D**) and von Frey (Student’s t-test, t = 1.886, ns = not significant) (**E**) tests showing that the conditional exclusion of NOP in noradrenergic neurons does not alter baseline nociception. Data represented as mean ± SEM.

**Supplementary Figure 7.**
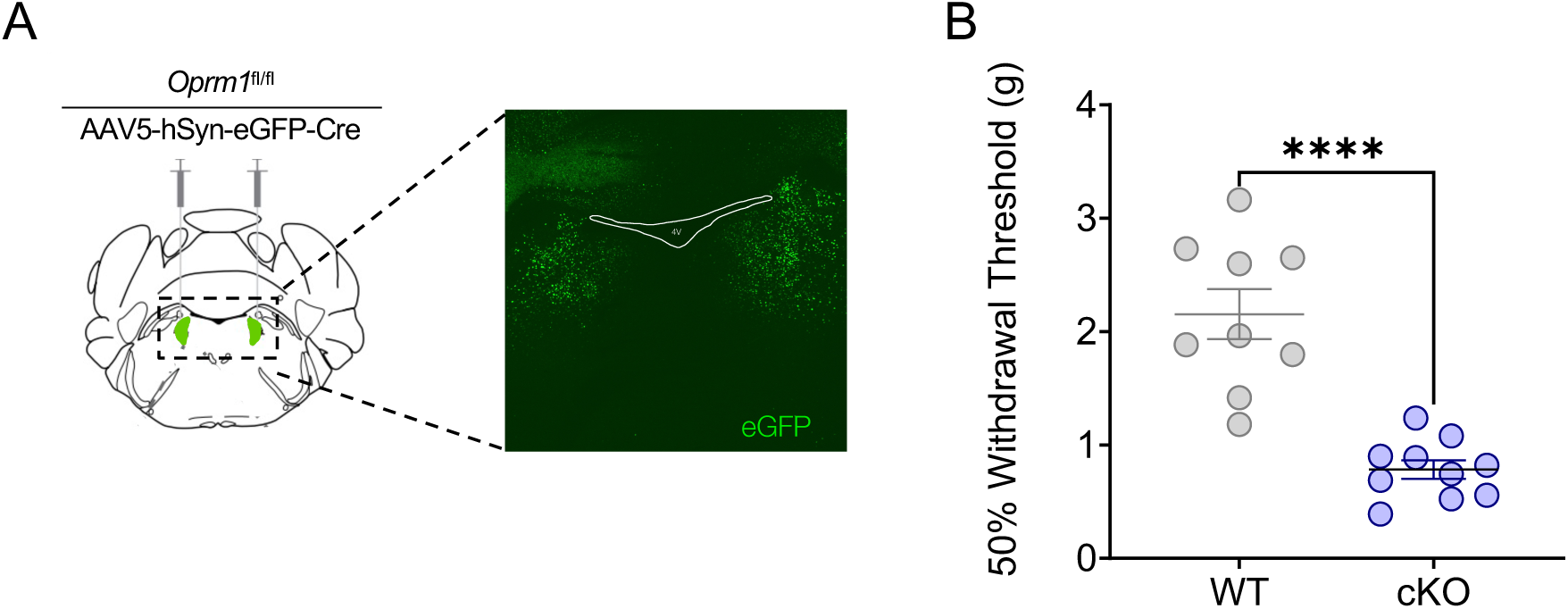
Spatial expression of Cre recombinase in the adult LC of O*prm1^fl/fl^*mice causes mechanical hypersensitivity. (**A**) Schematic and fluorescent image of the bilateral viral Cre expression in the LC. (**B**) *oprm1* deletion in the LC decreases baseline 50% mechanical withdrawal threshold. Student’s t-test, t = 6.06, ****p<0.0001. Data represented as mean ± SEM.

**Supplementary Figure 8.**
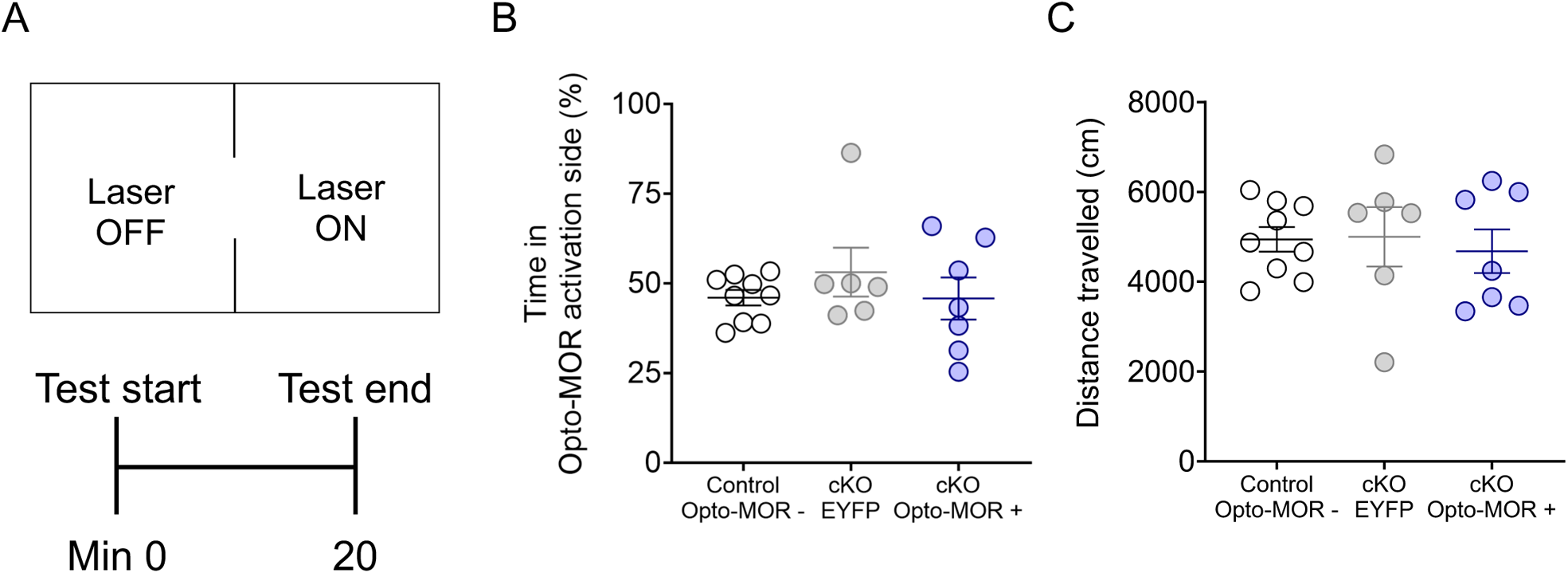
Optical activation of Opto-MOR signaling in LC of *Oprm1*^fl/fl^x*Dbh*^Cre+/−^ mice does not alter real-time place preference or locomotor behavior. (**A**) Schematics illustrating the experimental arrangement of real-time place preference test. (**B**) % time spent in Opto-MOR-paired side of real-time place testing apparatus shows no significant difference between groups. Kruskal-Wallis test, Kruskal-Wallis statistic = 0.4421. (**C**) Distance travelled during real-time place testing shows no significant effect of opto-MOR activation. One-way ANOVA, F = 0.1352. Data represented as mean ± SEM, no significant difference was found.

